# Systematic Identification of Microtubule Inner Proteins Reveals JPT2 as a Key Regulator of Lumen Microenvironment and Drug Sensitivity

**DOI:** 10.1101/2025.01.12.632577

**Authors:** Jinhui Shao, Rui Zhang, Peiyuan Wu, Honglin Xu, Zhengrong Zhou, Lusheng Gu, Lei Wang, Jinqi Ren, Xiahe Huang, Qi Xie, Yingchun Wang, Wei Ji, Wei Feng, Xin Liang, Wenxiang Meng

**Affiliations:** State Key Laboratory of Molecular Developmental Biology, Institute of Genetics and Developmental Biology, Chinese Academy of Sciences, Beijing 10019, China; University of Chinese Academy of Sciences, Beijing 100049, China; Neuroscience Center, Department of Basic Medical Sciences, Shantou University Medical College, Shantou, Guangdong 515041, China; National Laboratory of Biomacromolecules, CAS Center for Excellence in Biomacromolecules, Institute of Biophysics, Chinese Academy of Sciences, Beijing 100101, China; Wangjing Hospital of China Academy of Chinese Medical Sciences, Huajiadi Street, Chaoyang District, Beijing 100102, China; Bioland Laboratory (Guangzhou Regenerative Medicine and Health Guangdong Laboratory), Guangzhou, Guangdong, 510320, China; IDG/McGovern Institute for Brain Research, Tsinghua-Peking Joint Center for Life Science, School of Life Sciences, Tsinghua University, Beijing 100084, China; Innovation Academy for Seed Design, Chinese Academy of Sciences, 100101 Beijing, China

**Keywords:** Microtubule inner proteins, JPT2, MEC17, Paclitaxel

## Abstract

Microtubules are primarily studied for the interactions of proteins that bind to their outer surfaces and ends, while the regulatory mechanisms within the microtubule lumen, particularly in singlet microtubules critical for essential cellular processes, remain largely unexplored. Our study provides the first systematic identification of key regulatory proteins within the single microtubule lumen. Using proximity-dependent biotin identification (Bio-ID) coupled with mass spectrometry, we identified candidate microtubule inner proteins (MIPs), including Jupiter microtubule-associated homolog 2 (JPT2). JPT2 binds directly to microtubules and specifically localizes within the lumen, where it modulates the luminal environment by inhibiting acetylase MEC17 and independently affects the binding and efficacy of Paclitaxel. Furthermore, our screening identified additional MIPs that influence cellular sensitivity to Paclitaxel, indicating a link between luminal regulation and drug responsiveness. These discoveries reveal JPT2’s critical role in singlet microtubule regulation and suggest new therapeutic targets for enhancing cancer drug sensitivity.

## Introduction

Microtubules are a crucial component of the cell’s cytoskeleton, essential for maintaining cell structure, facilitating intracellular transport, and playing key roles in developmental diseases and cancer (1-4). While the interactions of proteins with the outer surfaces and ends of microtubules have been extensively studied, the regulatory mechanisms within the microtubule lumen remain largely unexplored, particularly in singlet microtubules, which are vital for essential cellular processes such as cell division, intracellular transport, and signal transduction (5-7). Understanding the luminal environment is crucial for revealing how microtubule dynamics and stability impact cellular behavior and drug responses.

The hollow lumen of microtubules, which can be up to 15-17 nm wide, contains post-translational modifications of tubulins, MIPs (8-10), and small molecules (11-14). These components collectively form the microtubule luminal environment, playing a key role in microtubule dynamics and cellular pharmacology. Studying how MIPs regulate this environment will provide insights into cellular functions and help develop new therapeutic approaches.

Since the 1960s, electron microscopy has revealed dense granules and protein particles within microtubules in various organisms (9, 15). More recently, cryo-electron tomography has uncovered periodic protein arrangements within the lumen of doublet microtubules, such as those found in the respiratory cilia of bovines and the flagella of human and mouse sperm (16, 17). However, despite singlet microtubules being more widely distributed and involved in diverse cellular processes, research into their lumenal regulation has lagged behind. Given their broader roles in cellular function, investigating the regulatory mechanisms of singlet microtubules is critical for a comprehensive understanding of microtubule biology.

Despite this, key findings have emerged: Tau and MAP6 proteins have been found localized to the inner walls of microtubules, where they stabilize microtubules and induce bending, respectively (18, 19). CSPP1, related to spindles and cilia, has also been shown to localize in the lumen of microtubules, contributing to their stabilization (20). Additionally, acetylation is the only known post-translational modification that occurs within the microtubule lumen, and studies have shown that the acetyltransferase MEC17/ATAT1 enters the microtubule lumen through its ends to promote microtubule acetylation (21-23).

To address the lack of knowledge regarding MIPs in singlet microtubules, we developed a novel screening method using proximity-dependent biotin identification (Bio-ID) coupled with mass spectrometry. This approach allowed us to systematically identify several candidate MIPs, including Jupiter microtubule-associated homolog 2 (JPT2). Our experiments revealed that JPT2 directly binds to microtubules and localizes within the lumen, modulating the luminal environment. Notably, JPT2 alters the positioning of the acetylase MEC17 and the chemotherapy drug Paclitaxel, influencing cellular resistance to Paclitaxel. Structural analysis using AlphaFold3 suggests that JPT2 may interact with β-tubulin at the Paclitaxel-binding site, further supporting its role as a key regulatory protein within the microtubule lumen.

In summary, our research introduces a novel approach for identifying MIPs, revealing their crucial roles in singlet microtubule regulation. Identifying JPT2 and other MIPs as modulators of drug sensitivity highlights their potential as therapeutic targets, particularly for enhancing the efficacy of cancer treatments like Paclitaxel.

## Results

### Establishment of the MIPs Identifying System

To comprehensively identify MIPs, we developed a protein capture system using proximity labeling combined with mass spectrometry analysis (Figure 1A). The N-terminus and C-terminus of β-tubulin are known to localize inside the microtubule lumen and outside, respectively (11, 24). For further study, we selected TUBB4B because it is highly expressed in the cervical cancer cell line HeLa cells and exists as a single isotype (25). We integrated the HA-tagged BirA* biotin ligase gene at both the start and stop codons of TUBB4B using the CRISPR-Cas9 system, successfully tagging both the N-terminus and C-terminus of the TUBB4B protein (26-28) (Supplementary Figure 1A-F).

**Figure 1.**
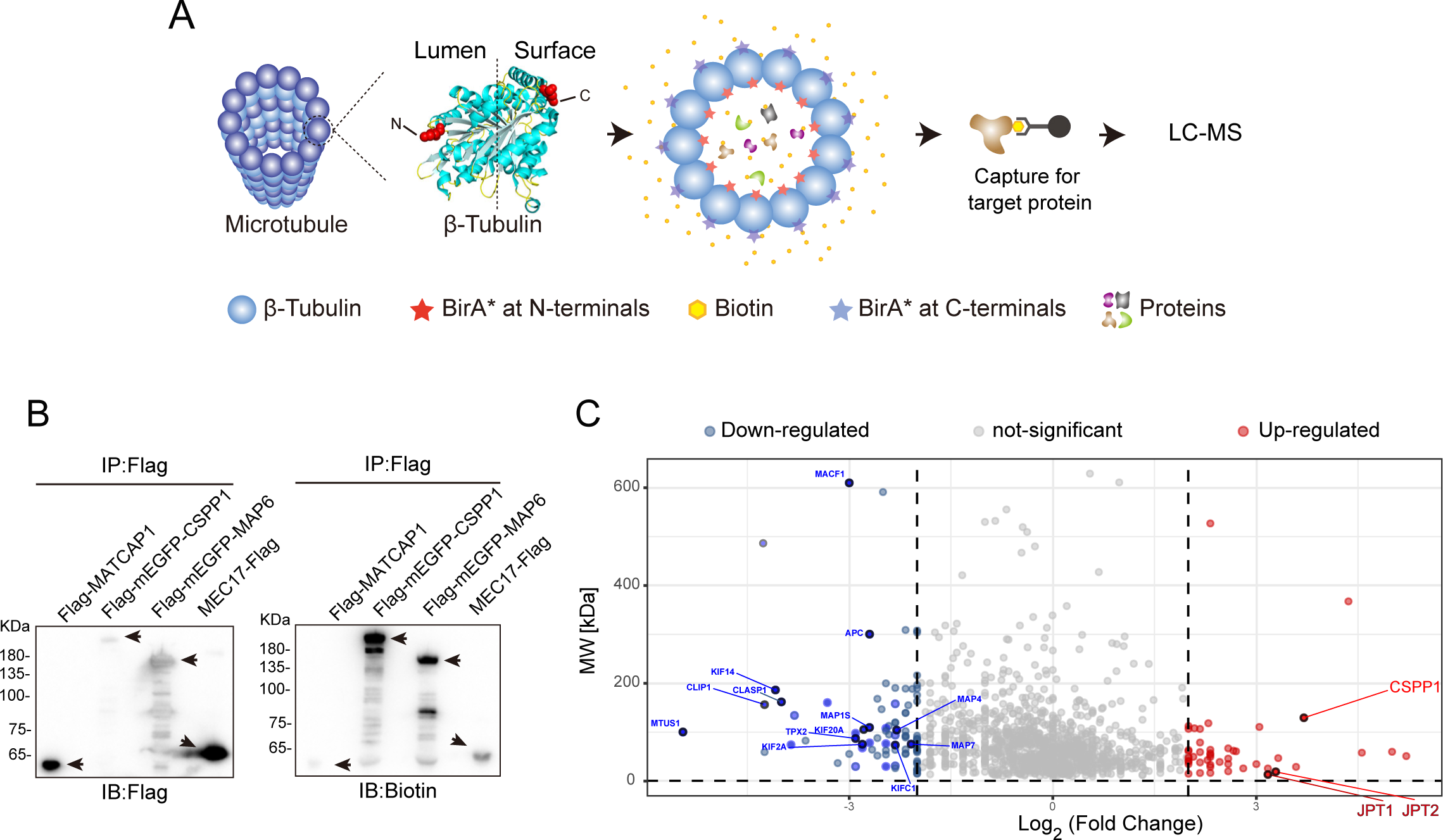
Establishment of the Singlet MIPs Screening System. **(A)** Schematic diagram of the MIP screen system. HA-BirA*-TUBB4B knock-in cells catalyzed biotinylation of the MIPs, then enrichment of biotinylated proteins using streptavidin-coated beads for isolation. **(B)** Overexpression of exogenous cytoplasm MIPs (CSPP1, MAP6, MEC17) and the microtubule surface protein MATCAP1 in HA-BirA*-TUBB4B knock-in cells, followed by immunoprecipitation using Flag tag to purify these proteins. Western blotting was then used to detect the biotin modification level and Flag expressed level on the same PVDF membrane. **(C)** Bioinformatic analysis of HA-BirA*-TUBB4B cells and TUBB4B-BirA*-HA cells proteomic data. The volcano plot’s vertical axis represents the proteins’ molecular weight, while the horizontal axis shows the Log_2_ ratio of HA-BirA*-TUBB4B (N-termini) to TUBB4B-BirA*-HA (C-termini). Compared to the TUBB4B-BirA*-HA (C-termini) cells, the red portion of the plot indicates proteins that are high abundance in the HA-BirA*-TUBB4B (N-termini) cells, and the blue portion indicates proteins that are low abundance in the HA-BirA*-TUBB4B (N-termini) cells.

To validate the functionality of this system, immunofluorescence staining confirmed that the engineered TUBB4B fusion proteins were correctly expressed in HeLa cells and formed microtubules (Supplementary Figure 1B, C, E, and F). Additionally, Western blot analysis using biotin detection demonstrated that the fused BirA* biotin ligase retained its enzymatic activity (Supplementary Figure 1C and F). Furthermore, we exogenously expressed HA-MATCAP1, which is known to localize to the outer surface of microtubules (29), alongside Flag-mEGFP-CSPP1, Flag-mEGFP-MAP6, and MEC17-mEGFP, which are located within the lumen of microtubules (19, 20, 23) (Supplementary Figure 1G). Proximity labeling experiments followed by immunoprecipitation using anti-Flag antibodies were then conducted (Figure 1B). The results revealed that CSPP1, MAP6, and MEC17 were effectively labeled with biotin, except for MATCAP1. These findings demonstrate that our MIPs identification system is robust and effective. It is important to note that cells must be permeabilized to remove free cytoplasmic signals before conducting immunofluorescence staining to observe the distribution of MIPs (30).

Using this system, we identified several potential MIP proteins (Figure 1C). When comparing proteins labeled with TUBB4B-BirA* at the N- and C-termini, we observed that MCAF1, CLASP1, MAP4, MAP7, and various KIF proteins - known to interact with the outer surface of microtubules (31-36) - were more abundant in proteins labeled by BirA* at the TUBB4B C-terminus. Conversely, proteins known to reside in the microtubule lumen, such as CSPP1 (20), were more enriched in proteins labeled with BirA* at the N-terminus. These results provided us with candidate proteins for potential MIPs.

### JPT2 is a MIP

We prioritized proteins with lower molecular weights, as most reported MIPs in singlet or doublet microtubules have molecular weights below 100 kD (16-20, 37-39) (Supplementary Figure 1 H). Among the identified proteins, JPT2 (also known as HN1L) attracted our attention (Figure 1C and 2A). JPT2 is a homolog of the Drosophila Jupiter protein (40-42). While JPT2 has been strongly linked to multiple cancers in human cells and to resistance against Paclitaxel treatment, its molecular mechanism remains poorly understood (43-49).

**Figure 2.**
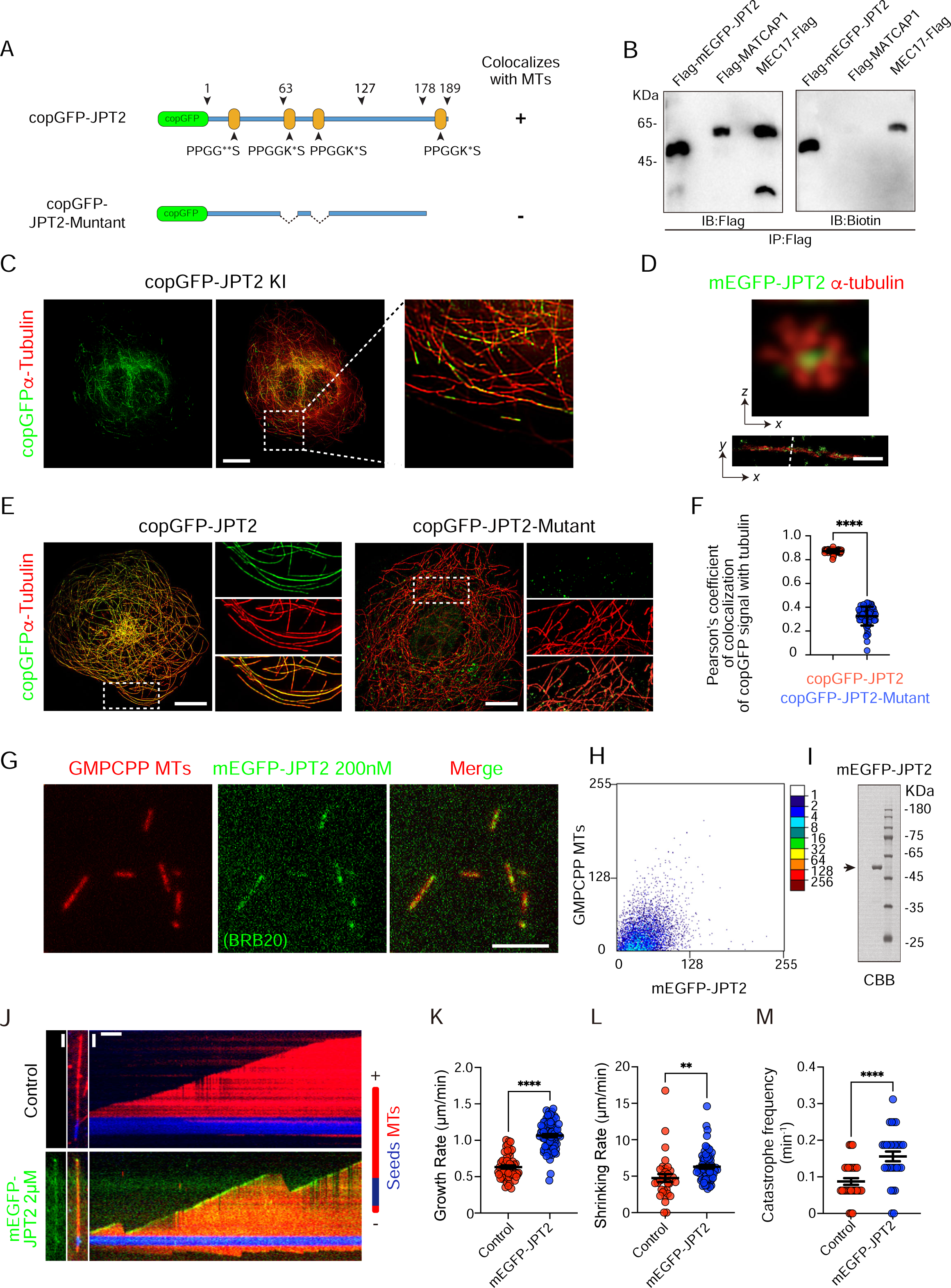
JPT2 is a singlet MIP. **(A)** Schematic illustration of the full-length JPT2 and the JPT2 mutant, which lacks microtubule binding capability. **(B)** Overexpressed MEC17 (MIP), JPT2, and MATCAP1 (outer MAP) in HA-BirA*-TUBB4B knock-in cells were purified using immunoprecipitation with the Flag tag. Western blotting was then used to detect the biotin modification and Flag expression levels on the same PVDF membrane. **(C)** Immunofluorescence staining of α-tubulin (red) in cop-GFP-JPT2 knock-in Hela cells, scale bars: 10 μm.**(D)** 3D single-molecule localization microscopy (SMLM) image of mEGFP-JPT2 (green) and α-tubulin (red) in HeLa cells, scale bars: 2 μm **(E)** Immunofluorescence staining of α-tubulin (red) in copGFP-JPT2 and copGFP-JPT2-mutant expressing HeLa cells, scale bars: 10 μm. **(F)** Quantification of the localization of copGFP-JPT2 and copGFP-JPT2-mutant with α-tubulin. Each symbol represents an individual cell. For copGFP-JPT2, n = 67 cells; for copGFP-JPT2-mutant, n = 63 cells. **(G)** TIRF microscopy images showing colocalization of strep-mEGFP-JPT2 (green) with GMPCPP microtubule seeds (Red). Scale Bar: 5 μm. **(H)** Co-localization of mEGFP-JPT2 and GMPCPP microtubules were analyzed using the scatterJ plug-in in ImageJ. The horizontal and vertical axes of the scatterplot represent the gray values obtained for each pixel point in each channel, and the closer the image is to the diagonal, the higher the degree of co-localization is indicated. **(I)** Coomassie Brilliant Blue staining of a gel with mEGFP-JPT2 purified from HEK293 cells. **(J)** Representative kymographs of control and mEGFP-JPT2 binding microtubules. Scale bars: vertical, 2 μm; horizontal, 1 min. **(K)** Quantification of the microtubule plus end growth rate binding with mEGFP-JPT2 proteins or control, n = 58 for control group; n = 80 for the mEGFP-JPT2 group. **(L)** Quantification of the shrinking rate of microtubule plus ends binding with mEGFP-JPT2 proteins or control, n = 33 for the control group; n = 64 for the mEGFP-JPT2 group. **(M)** Quantification of the catastrophe frequency of microtubule plus ends binding with mEGFP-JPT2 proteins or control, n = 30 for control and mEGFP-JPT2 group. Values = means ± SEM. unpaired two-tailed Student’s t-test, ****p < 0.0001.

To confirm these findings, we transfected the HA-BirA*-TUBB4B (N-termini) knocked-in HeLa cell line with MEC17-Flag, Flag-MATCAP1, and Flag-mEGFP-JPT2, respectively, and conducted proximity labeling experiments (Figure 2B). Notably, the lumenal BirA* specifically biotinylated JPT2 and MEC17, but not MATCAP1, indicating that JPT2 is a novel MIP candidate.

To investigate the localization of JPT2 within cells, we faced the challenge of the lack of suitable specific antibodies for immunofluorescence staining of endogenous JPT2. Therefore, we used CRISPR/Cas9 to integrate a copGFP tag at the initiation codon of the JPT2 gene (copGFP-JPT2) in HeLa cells (Figure 2A and 2C). Immunofluorescence staining with anti-α-tubulin antibodies revealed colocalized copGFP and α-tubulin signals, suggesting that JPT2 localizes to microtubules at endogenous levels. Notably, JPT2’s distribution along microtubules was discontinuous (Figure 2C).

Furthermore, to confirm JPT2’s localization within the microtubule lumen and to intensify the signal, we transfected mEGFP-JPT2 into HeLa cells and performed immunostaining (Figure 2D). Super-resolution microscopy (ROSE-Z)(50), which allows visualization of the microtubule lumen due to its high axial resolution, revealed that the mEGFP signal was localized within the microtubule lumen, enclosed by α-tubulin, further supporting the luminal localization of JPT2.

### PPGGK*S Motif is Important for JPT2 Localization at the Microtubule

To further investigate the key sequences responsible for JPT2’s localization to microtubules, a series of JPT2 mutants were generated for screening (Supplementary Figure 2A). We expressed full-length JPT2 and its mutants, followed by immunostaining using anti-α-tubulin antibodies (Supplementary Figure 2, Figure 2 E and F). The results indicate that both amino acids 61^th^-80^th^, 81^th^-100^th^ and 178^th^-190^th^ can localize to microtubules, suggesting that JPT2 has multiple regions associated with its microtubule localization. Interestingly, JPT2 possesses four incompletely repeated conserved sequences, which include the PPGG amino acid residues. Notably, the last three of these sequences are located in regions critical for microtubule localization (Supplementary Figure 2A and B).

We found that the characteristic sequence within the PPGG sequence of JPT2 is similar to the PGGG motif, a sequence found in the Tau protein that plays a key role in its localization to the microtubule lumen(51). Further experiments found that mutants lacking the last three incompletely repeated conserved PPGG sequence groups completely lost the ability to localize to microtubules (Figure 2 E and F). Compared to the first incompletely repeated conserved sequences containing the PPGG sequence, these motifs have an additional lysine (K) after the PPGG sequence (Figure 2A). These results suggest that the PPGGK*S motif may play an important role in JPT2 localization.

### JPT2 Directly Binds to Microtubules

To further investigate the interaction between JPT2 and microtubules, we purified JPT2 and mEGFP-JPT2 using the pHUE expression system in bacteria and the Strep-tagII expression system in HEK293 cells, respectively. Co-sedimentation assays confirmed JPT2’s ability to interact directly with microtubules (Supplementary Figure 3A-C). Total internal reflection fluorescence (TIRF) microscopy showed that mEGFP signals overlapped with GMPCPP-stabilized microtubules (Figure 2G-I), providing further evidence that JPT2 binds directly to microtubules.

We performed an in vitro microtubule polymerization assay and observed the process in detail using TIRF microscopy (Figure 2J). Fluorescence signals of mEGFP-JPT2 were detected along the microtubules, with a notably stronger intensity at the microtubule plus end. This observation suggests that JPT2 may influence microtubule dynamics. Statistical analysis revealed that JPT2 significantly increased both the growth rate and the retraction rate at the microtubule plus end (Figure 2K-M), indicating that JPT2 plays a critical role in regulating microtubule dynamics.

Our findings were further validated in vivo. We transfected copGFP-IRES-EB3-mScarlet and copGFP-JPT2-IRES-EB3-mScarlet into HeLa cells to study the microtubule plus end dynamics (Supplementary Figure 3D). Live-cell imaging demonstrated a significant increase in the growth rate of EB3-labeled growing microtubule ends in cells expressing JPT2, along with a shortened life cycle (Supplementary Figure 3E). These results are consistent with our in vitro data on microtubule dynamics (Figure 2K-M), emphasizing JPT2’s direct and significant role in precisely controlling microtubule dynamics.

### JPT2 Prevents MEC17 Microtubule Localization

The 15-17 nm lumen of microtubules constitutes a microenvironment that includes various luminal proteins, amino acids of tubulin exposed to the lumen, and small molecules bound to the microtubule inner surface (14, 19, 20). Since K40 acetylation of α-tubulin occurs on the inner surface of microtubules (52, 53), and JPT1/HN1 has been shown to affect this modification (54). We explored whether these microtubule luminal environmental factors might be associated with each other.

First, we conducted immunostaining using knocked-in copGFP-JPT2 HeLa cells to investigate the potential relationship between JPT2 and microtubule acetylation modification (Figure 3A -C). We found that the colocalization rate of JPT2 with acetylated microtubules was significantly lower than that with normal microtubules (Figure 2B), suggesting that there may be a potential relationship between JPT2 and microtubule acetylation modification. However, increasing the acetylation level of microtubules by adding TSA (microtubule deacetylase inhibitor) did not change the distribution of JPT2 on microtubules (Supplementary Figure 4 A). Therefore, we concluded that the K40 acetylation site of α-tubulin is not directly related to the localization of JPT2. Next, we studied the impact of overexpressing and knocking out JPT2 on microtubule acetylation in HeLa cells (Supplementary Figure 4 B and C). Our findings indicate that overexpression of JPT2 leads to a decrease in microtubule acetylation level (Figure 3 D-F), while reducing its expression level does not affect microtubule acetylation level (Supplementary Figure 4 D and E). These results confirm our hypothesis that JPT2 may not be directly involved in regulating microtubule acetylation.

**Figure 3.**
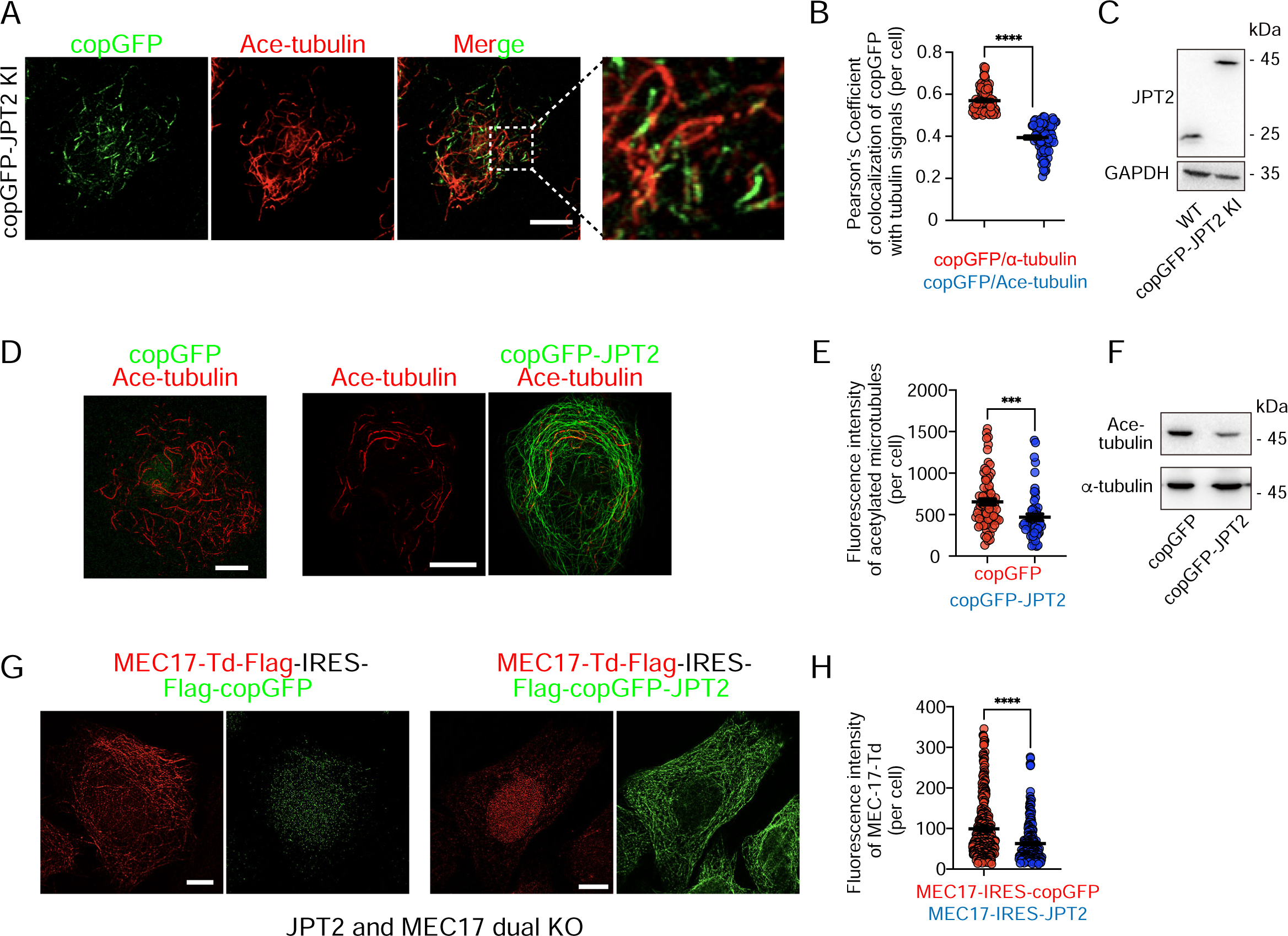
JPT2 inhibits microtubule acetylation by restricting MEC17 binding to microtubules. **(A)** Immunostaining for Ace-tubulin (red) in copGFP-JPT2 knock-in HeLa cells, scale bars: 10 μm. **(B)** Quantification of the colocalization of copGFP-JPT2 and Ace-tubulin or α-tubulin. Each symbol represents an individual cell. For the α-tubulin group, n = 101 cells; for the Ace-tubulin group, n = 88 cells. **(C)** Western blotting was used to detect JPT2 expression levels in wild-type HeLa cells and copGFP-JPT2 knock-in HeLa cells. GAPDH as a reference protein. **(D)** Immunostaining for Ace-tubulin (red) in copGFP-JPT2 and copGFP stable expressing HeLa cells, scale bars: 10 μm. **(E)** Quantification of the mean Ace-tubulin intensity in copGFP-JPT2 and copGFP stable expressing HeLa cells. The mean intensity of cells overexpressing copGFP-JPT2 was normalized to the mean intensity in control cells. Total number of cells analyzed: control cells, n = 77; cells overexpressing copGFP-JPT2, n = 55. Bars represent pooled data from two independent experiments. **(F)** Western blotting was used to detect Ace-tubulin expression level, α-tubulin as a reference protein. **(G)** TD-tomato (red) and CopGFP (Green) signals were obtained in MEC17-TD-tomato-Flag-IRES-Flag-copGFP and MEC17-TD-tomato-Flag-IRES-Flag-copGFP-JPT2 expressing HeLa cells respectively, scale bars: 10 μm. **(H)** Quantification of the mean MEC17 intensity from images as in (G). The total number of cells analyzed: MEC17-TD-tomato-OE cells, n = 349; cells overexpressing MEC17-TD-tomato-IRES-CopGFP-JPT2 cells, n = 237. Bars represent pooled data from two independent experiments. Values = means ± SEM. ***p < 0.001, ****p < 0.0001; unpaired two-tailed Student’s t-test.

Given that microtubule acetylation and deacetylation are dynamically regulated by specific enzymes (21, 55, 56), we explored the relationship between JPT2 and microtubule deacetylases and acetylases. First, we investigated the possibility that JPT2 regulates microtubule acetylation by interacting with microtubule deacetylases. Immunoprecipitation experiments showed that JPT2 (HA-BirA*-JPT2) did not interact with microtubule deacetylases HDAC-6 or SIRT-2 (Supplementary Figure 4 F), demonstrating that the relationship between the distribution of JPT2 on microtubules and deacetylated microtubules was not the result of direct regulation by microtubule deacetylases.

Does JPT2 disrupt the normal function of microtubule acetyltransferase? To exclude the influence of microtubule background acetylation, we established a HeLa cell line with double knockout of MEC17 and JPT2 genes (Supplementary Figure 4 G and H) and then transfected MEC17-tdTomato-Flag-IRES-Flag-copGFP and MEC17-tdTomato-Flag-IRES-Flag-copGFP-JPT2 into the above cells (Figure 3 G and H). Immunofluorescence staining results showed that in cells co-expressing copGFP, MEC17 was colocalized with microtubules. However, in cells co-expressing JPT2, MEC17 lost its microtubule localization, suggesting that JPT2 prevents MEC17 microtubule localization and potentially impacts the Microtubule luminal environment.

The N- and C-termini of Jupiter in Drosophila, and JPT1 and JPT2 in mammals are highly conserved (Supplementary Figure 5 A). We transfected copGFP-Jupiter into HeLa cells and observed that Jupiter showed strong signals in the nucleus and co-localized with microtubules (Supplementary Figure 5 B). When the 17-amino acid sequence, which includes an incomplete repeat of the conserved PPGGK*S sequence at Jupiter’s C-terminus, was overexpressed in HeLa cells (Supplementary Figure 5 C), it was unexpected to see a notable alteration in microtubule acetylation, with the continuous signal transforming into a dot (Supplementary Figure 5 C-E). This suggests that influence on the microtubule lumen environment is a conserved function among Jupiter family proteins.

### JPT2 Competes with Paclitaxel for Microtubule Binding Sites

Small molecules bound to the inner surface of microtubules are also important elements of the microtubule environment (14). Based on the known association of the PGGG motif with the β-tubulin Paclitaxel binding site (18), we used AlphaFold3 to predict how JPT2 interacts with microtubules (Figure 4 A and B). Our simulations focused on the 13 amino acids at the C-terminus of JPT2 (Figure 4 A), as our experiments demonstrated the presence of the PPGGK*S motif and significant localization with microtubules (Supplementary Figure 2). The results showed that the series at positions 186 and 188 of the 13 amino acids (LNPPGGKSSISFY) at the C-terminus of JPT2 could bind to aspartic acid at position 224 (Asp224) and arginine (Arg) at position 276 of the core helix H7 and M-loop of β-tubulin (57), respectively (Figure 4B). When we mutated these two key serines to alanine, the co-localization phenomenon between the C-terminus of JPT2 and the microtubules disappeared (Figure 4C and D). This result is consistent with the AlphaFold3 prediction.

**Figure 4.**
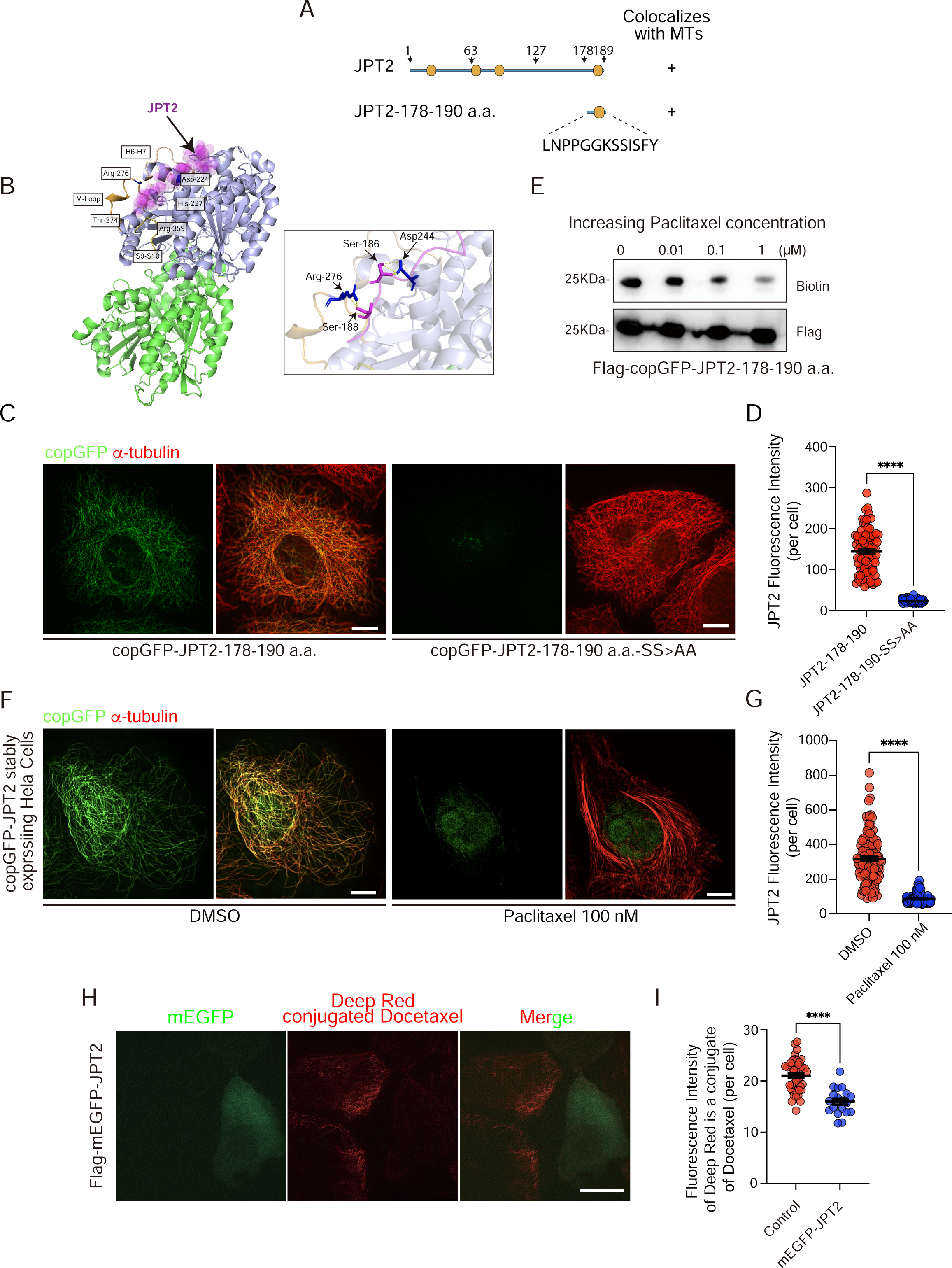
Paclitaxel disassociates JPT2 from microtubules. **(A)** Schemes of the domain organization of the full-length JPT2, JPT2-178-190 a.a. which contains the minimal microtubule-binding motif of JPT2. **(B)** The alpha-fold prediction suggests that the binding site of JPT2-178-190 a.a. with β-tubulin is similar to the binding site of Paclitaxel. **(C)** Immunostaining for α-tubulin (red) in copGFP-JPT2-178-190 a.a. and copGFP-JPT2-178-190 a.a.-SS > AA expressing HeLa cells were shown in the right panel, scale bars: 10 μm. **(D)** Quantification of the copGFP-JPT2-178-190 a.a. and copGFP-JPT2-178-190 a.a.-SS > AA intensity in HeLa cells. For the copGFP-JPT2-178-190 a.a. group, n = 83 cells; for the copGFP-JPT2-178-190 a.a.-SS > AA group, n = 79 cells; Bars represent pooled data from two independent experiments. **(E)** Flag-copGFP-JPT2-178-190 a.a. overexpressed in the HA-BirA*-TUBB4B knock-in cells, catalyzed biotinylation and treated with increasing concentrations Paclitaxel, followed by IP purify Flag-copGFP-JPT2-178-190 a.a. protein, Western blotting detected proteins Biotin modification level and Flag expressed level in same PVDF-membrane. **(F)** Immunostaining for α-tubulin (red) in DMSO and Paclitaxel-treated copGFP-JPT2 stably expressing HeLa cells, scale bars: 10 μm. **(G)** Quantification of the JPT2 intensity in DMSO and Paclitaxel-treated HeLa cells. The mean intensity of cells treated with Paclitaxel was normalized to that in cells treated with DMSO. Total number of cells analyzed: control cells, n = 111; Paclitaxel group, n = 118. Bars represent pooled data from two independent experiments. **(H)** Time-lapse images of Deep Red conjugated Docetaxel (red) binding microtubule experiment in mEGFP-JPT2 overexpressed HeLa cells, scale bars: 10 μm. **(I)** Quantification of the mean Docetaxel intensity in JPT2 overexpressed cells and un-transfected cells. The mean intensity of cells overexpressing mEGFP-JPT2 was normalized to the mean intensity in control cells. Total number of cells analyzed: control cells, n = 44; Paclitaxel group, n = 19. Bars represent pooled data from two independent experiments. Values = means ± SEM. ****p < 0.0001; unpaired two-tailed Student’s t-test.

This allowed us to investigate the impact of JPT2 and Paclitaxel on the binding of Asp224 and Arg227, which are key amino acids for Paclitaxel binding (57). We transfected HA-BirA*-TUBB4B knock-in HeLa cells with Flag-copGFP-JPT2 (178-190 a.a.) and conducted proximity labeling assays using different concentrations of Paclitaxel (Figure 4E). We observed a decrease in the biotinylation of Flag-copGFP-JPT2 (178-190 a.a.) with increasing Paclitaxel concentrations, indicating that Paclitaxel affects the biotinylation of JPT2’s C-terminus. Additionally, we treated HeLa cells stably expressing full-length copGFP-JPT2 and Flag-copGFP-JPT2 (178-190 a.a.) with 100 nM Paclitaxel for 12 hours and performed immunostaining, which revealed a reduction in the microtubule localization of JPT2 upon Paclitaxel administration (Figure 4 F and G, Supplementary Figure 6 A and B). These results suggest that Paclitaxel may affect the microtubule lumenal distribution of JPT2 within the cell. It is important to note that the microtubule localization of MAP6, typically found in the microtubule lumen, remained unaffected by Paclitaxel (Supplementary Figure 6 C and D).

To investigate whether microtubule stabilization was responsible for its disruption, we treated the copGFP-JPT2 stably expressed HeLa cells with Paclitaxel and other non-taxane stabilizers interacting with different binding sites (Supplementary Figure 7 A and B). The results showed that treatment with Paclitaxel, epothilone D (Epo D), and TPI287 disrupted JPT2 microtubule localization, whereas noscapine and TTI237 did not affect its localization (Supplementary Figure 7 C and D). Thus, JPT2’s disrupted microtubule attachment is specifically caused by taxanes, not by an overall change in microtubule stability.

Lastly, HeLa cells overexpressing Flag-mEGFP-JPT2 treated with low concentrations of deep red fluorophore-conjugated Docetaxel showed reduced Docetaxel binding to microtubules in cells with high JPT2 expression by time-lapse recording (Figure 4H and I). Our results reveal that JPT2 and Paclitaxel mutually influence their localization within the microtubule lumen, suggesting JPT2 as a key modulator of Paclitaxel efficacy and a potential target for overcoming drug resistance.

### Paclitaxel may change the microtubule luminal environment

Our results suggest that the binding of Paclitaxel to microtubules could alter the microtubule luminal environment (Figure 4F and G). Conversely, do these small molecules affect the composition of proteins within the microtubule lumen?

To screen for proteins that respond to Paclitaxel (Figure 5A). After gradient administration of Paclitaxel to cells, we performed proximity labeling and mass spectrometry identification and used the MIP screening system to identify 29 proteins sensitive to Paclitaxel (Figure 5B). Interestingly, CSPP1, which is currently reported to be localized in the microtubule lumen (20), JPT2 and JPT1, which we found are all among them. This result supports our conclusion that JPT2 competes with Paclitaxel for binding sites in the microtubule lumen and coincides with the hypothesis that Paclitaxel may affect the lumen environment more broadly.

**Figure 5.**
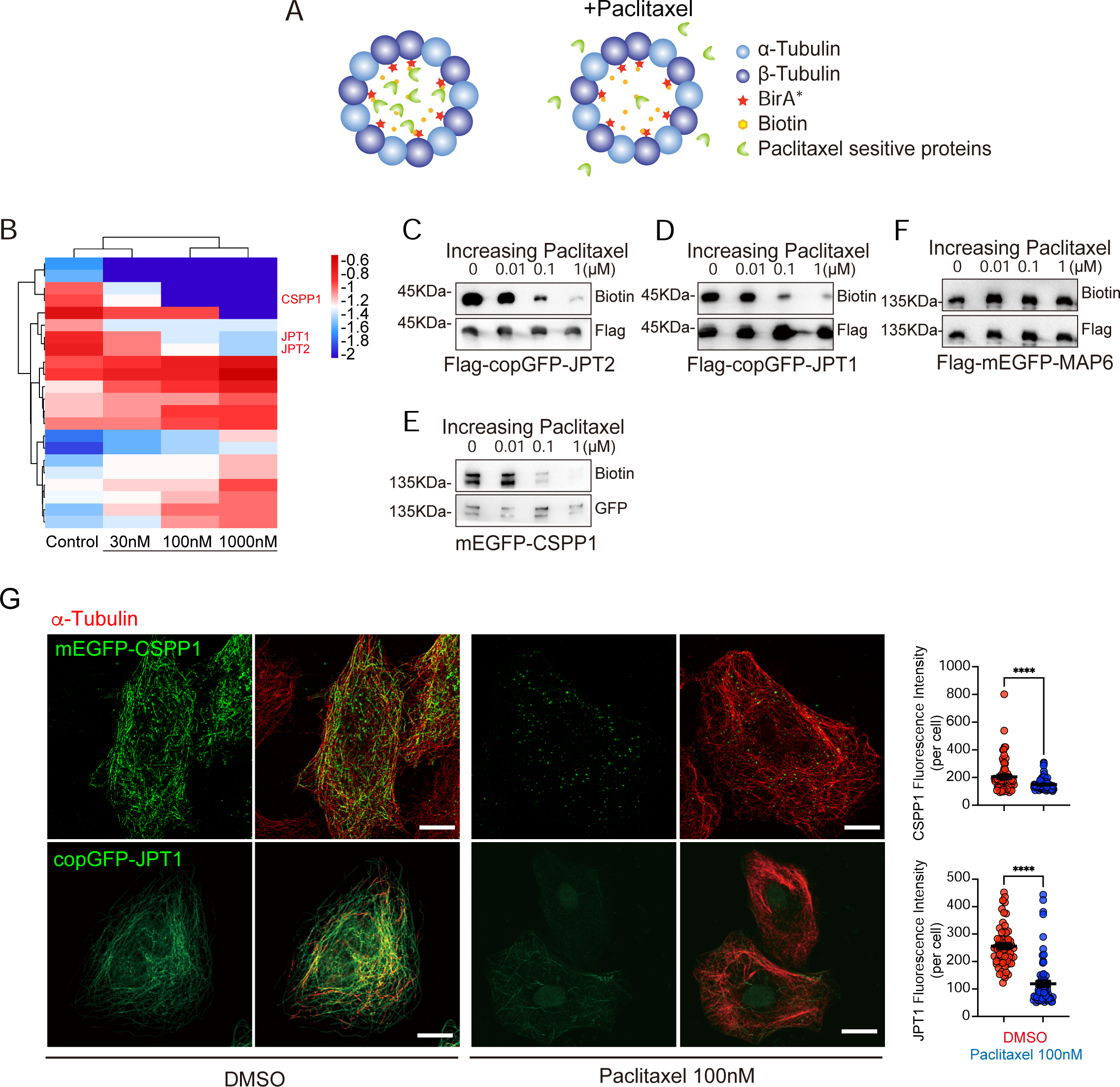
JPT1 and CSPP1 were new Paclitaxel-sensitive MIPs. **(A)** Schematic diagram showed HA-BirA*-TUBB4B knock-in cells undergoing biotinylation of MIPs and treatment with increasing concentrations of Paclitaxel, followed by streptavidin bead enrichment to capture the proteins. **(B)** Bioinformatic analysis of proteomic data found 3 new Paclitaxel-sensitive MIPs. The heat map shows the changes in MIPs in different concentrations of Paclitaxel treatment groups. **(C-F)** Falg-copGFP-JPT2 **(C),** Falg-copGFP-JPT1 **(D)**, mEGFP-CSPP1 **(E)** and Falg-mEGFP-MAP6 **(F)** overexpressed in the HA-BirA*-TUBB4B knock-in cells, catalyzed biotinylation in the MIP and treated with increasing concentrations Paclitaxel, followed by IP purify MIP, Western blotting detected proteins Biotin modification level and Flag/GFP expressed level in same PVDF-membrane. **(G)** Immunostaining for mEGFP-CSPP1, copGFP-JPT1 and α-tubulin (red) in DMSO and Paclitaxel-treated HeLa cells, scale bars: 10 μm. Quantification of the CSPP1 and JPT1 intensity in DMSO and Paclitaxel-treated HeLa cells are shown in the right panel. For the CSPP1 DMSO group, n = 70 cells; for the CSPP1 Paclitaxel group, n = 100 cells. For the JPT1 DMSO group, n = 64 cells; for the JPT1 Paclitaxel group, n = 76 cells. Values = means ± SEM. ****p < 0.0001, unpaired two-tailed Student’s t-test.

Therefore, we used biochemistry and cell biology methods to verify the above results. In this experiment, we used MAP6 (19), which is currently reported to work in the microtubule lumen but is not included in our candidate proteins (Figure 5B). We found that after administration of Paclitaxel to cells, the biotinylation of CSPP1, JPT1 and JPT2 was significantly reduced (Figure 5 C-E), while the biotin labeling of MAP6 did not change (Figure 5F). Immunofluorescence staining also obtained the same results. The localization of CSPP1, JPT1 and JPT2 to microtubules all responded to the addition of paclitaxel (Figure. 5G, Figure. 4F and G), whereas MAP6 did not change (Supplementary Fig. 6C and D). Furthermore, time-lapse recordings show that overexpression of these proteins can reduce the binding of deep red fluorophore-conjugated docetaxel to microtubules in cells (Supplementary Figure 8A), like JPT2 (Figure 4H and I). Our results indicate that these MIPs and Paclitaxel can affect each other’s microtubule localization.

### Paclitaxel-sensitive MIPs are Associated with Paclitaxel Resistance

Paclitaxel is a widely used cancer treatment that targets microtubules to disrupt their stability and dynamics, blocking cell division and causing apoptosis. However, resistance to Paclitaxel remains a significant clinical challenge (58-60).

Our data indicate that paclitaxel-sensitive MIPs also affect the interaction between paclitaxel and microtubules. First, we used the TCGA database to study the relationship between the expression levels of these genes and docetaxel, and we found that high expression of the above proteins was associated with reduced efficacy of docetaxel in a variety of cancers (Supplementary Figure 8B). At the same time, it has been reported that reducing the expression level of JPT2 in docetaxel-resistant ESCC cells can increase their sensitivity to docetaxel (48). These findings establish Paclitaxel-sensitive MIPs, including JPT2, as critical regulators of drug resistance, positioning them as promising targets for overcoming chemotherapy resistance.

Next, we focused on JPT2. To ask whether the microtubule lumenal localization of JPT2 is associated with the Paclitaxel resistance. We utilized a GAPDH promoter-driven gene expression knock-in system to ensure sustained and stable expression of the target gene while minimizing expression differences between vectors (Figure 6A) (61). We established GAPDH-P2A-HA-BirA*, GAPDH-P2A-HA-BirA*-JPT2 and GAPDH-P2A-HA-BirA*-JPT2 Mutant gene knock-in HeLa cell lines (Figure 6 B and C), and then evaluated cell viability using CCK8 assay after Paclitaxel treatment (Figure 6D). The results showed that cells with full-length JPT2 were significantly less sensitive to Paclitaxel than control cells, which was not reflected in cells with a JPT2 mutant lacking microtubule-binding ability. This result supports our speculation that the binding of JPT2 to microtubules is related to the sensitivity of cells to Paclitaxel.

**Figure 6.**
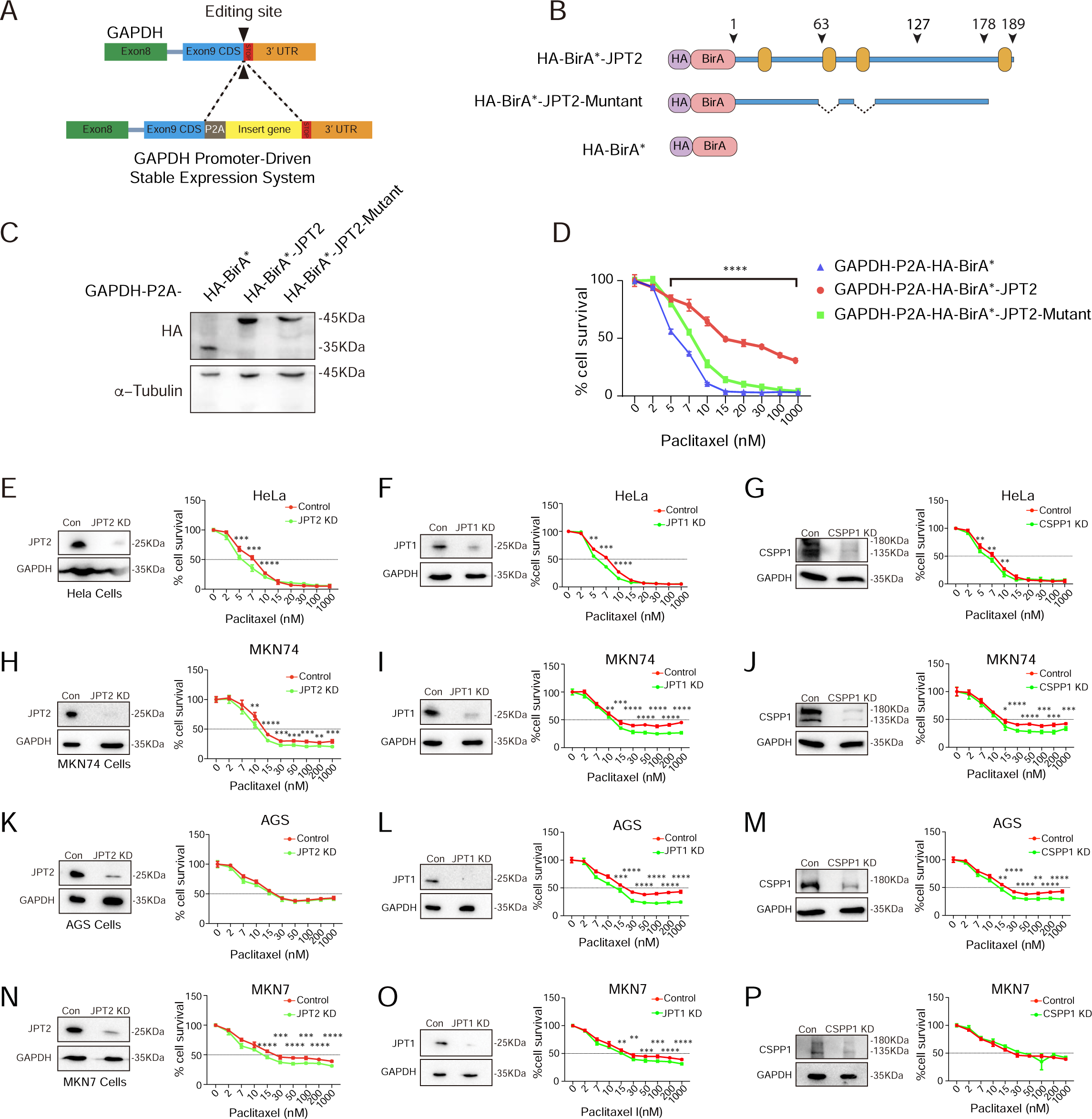
JPT2 is a Paclitaxel-sensitive protein and is associated with the Paclitaxel resistance of tumor cells. **(A)** Schematic diagram of GAPDH promoter-driven stable expression system. **(B)** Schemes of the domain organization of the HA-BirA*-JPT2, the HA-BirA*-JPT2-Mutant which lacks microtubule binding capability, and HA-BirA* as negative control. **(C)** Western blotting detected the expressing level of GAPDH-P2A-HA-BirA*, GAPDH-P2A-HA-BirA*-JPT2 and GAPDH-P2A-HA-BirA*-JPT2-mutant knock-in cells, α-tubulin as a reference protein. **(D)** Cell viability experiment of GAPDH-P2A-HA-BirA* knock-in cells, GAPDH-P2A-HA-BirA*-JPT2 knock-in cells and GAPDH-P2A-HA-BirA*-JPT2-mutant knock-in cells after 72 h of Paclitaxel treatment (increasing concentrations 0 to 1 μM; graphic representation of one experiment; lines show nonlinear curve fittings; error bars in each concentration show ± SD of five replicates). **(E-P)** Knockdown of JPT2, JPT1, and CSPP1 in HeLa cells **(E-G)**, MKN74 cells **(H-J)**, AGS cells **(K-M)**, and MKN7 cells **(N-P)**, with subsequent detection of cell viability in knockdown cells. After 72 hours of Paclitaxel treatment at increasing concentrations from 0 to 1 μM, a graphical representation of one experiment is shown. The lines represent nonlinear curve fittings, and the error bars in each concentration indicate ± SD from five replicates. Statistical analyses were performed with the Student’s t-test. *p < 0.05; **p < 0.01; ***p < 0.001; ****p < 0.0001.

Finally, to validate our findings, we knocked down JPT2 as well as CSPP1 and JPT1 in HeLa cells, and Paclitaxel-resistant MKN-74, MKN7, and AGS cells to test whether reducing the protein levels of these MIPs could improve their tolerance to Paclitaxel (Figure 6 E-P). Surprisingly, we found that knocking down these proteins increased the sensitivity of cells to Paclitaxel to varying degrees, although the extent varied between different cells.

## Discussion

In this study, we systematically identified key MIPs using a novel screening method. We uncovered that Jupiter microtubule-associated homolog 2 (JPT2) is pivotal in regulating the microtubule lumenal environment. Our findings demonstrate that JPT2 not only localizes within the microtubule lumen but also modulates both MEC17 acetylation and Paclitaxel binding. These results highlight the significance of microtubule lumenal proteins in controlling microtubule stability and drug efficacy, providing new insights into how these proteins affect cellular processes and chemotherapy outcomes.

### Microtubule Luminal Localization of JPT2

JPT2 was identified as one of the MIP through our system. JPT2 is a homolog of Drosophila’s microtubule-associated protein Jupiter and is highly associated with cancer progression and prognosis, though its interaction with microtubules is less studied (40, 62). We confirmed that JPT2 is a direct microtubule-binding protein, with super-resolution microscopy verifying its localization within the microtubule lumen.

We found that JPT2 has four imperfect repeat sequences, predominantly PPGG. Except for the N-terminal repeat, the three repeats starting from JPT2’s C-terminal 13 amino acids can localize to microtubules and exhibit the conserved PPGGK*S sequence. We observed that the PPGGK*S sequence bears a high similarity to the PGG motif, which plays a crucial role in the Tau protein, especially in the Paclitaxel binding pocket of β-tubulin. PGGG motif includes proline (P) and glycine (G), where proline induces specific folding and glycine provides flexibility. This combination forms a hairpin structure that fits well into the binding pocket, stabilizing microtubules and promoting their assembly (18). Using AlphaFold3, we predicted the interaction of JPT2’s C-terminal 13 amino acids containing PPGGK*S with α/β-tubulin dimer, suggesting it can bind the Paclitaxel binding pocket of β-tubulin similarly to the PGGG motif. This finding supports our observation that JPT2 and Paclitaxel mutually influence microtubule localization. We hypothesize that the additional proline in PPGGK*S enhances structural rigidity, while glycine retains the necessary flexibility. This combination of rigidity and flexibility is crucial for fitting the complex three-dimensional structure of the binding pocket. Our results indicate that PGGG and PPGGKS sequences have significant potential in developing microtubule-based nanomaterials and drug delivery systems.

### JPT2 regulates the Microtubule luminal environment

Microtubule acetylation is a biological event occurring within the microtubule lumen (63, 64). As a luminal protein, JPT2 reduces MEC17 localization on microtubules, thereby lowering microtubule acetylation levels in cells. However, even when MEC17 loses microtubule localization, microtubules still maintain a certain level of acetylation (65). We hypothesize that MEC17’s ability to acetylate microtubules is temporary, meaning stable localization of MEC17 on microtubules is unnecessary, which aligns with previous understandings. Another possibility is that MEC17 may activate other downstream acetyltransferases, which then localize to microtubules to complete acetylation. These hypotheses need further experimental validation.

Furthermore, the microtubule lumen environment is a crucial target for various small molecules and drugs (14). Paclitaxel, a common cancer chemotherapy drug, binds to the luminal part of β-tubulin, altering the conformation of the microtubule lattice to increase microtubule stability (66, 67). Our research shows that JPT2 and Paclitaxel mutually influence each other’s localization on microtubules. Additionally, we identified several potential MIPs sensitive to Paclitaxel using our MIP capture system. These results suggest that various factors within the microtubule lumen environment can interact with each other.

Overall, JPT2’s role in the microtubule lumen environment is multifaceted. It influences microtubule structure and function directly by binding to microtubules and indirectly by regulating other components and physicochemical properties of the lumen. Further research into JPT2’s role in the microtubule lumen environment will help us gain a more comprehensive understanding of microtubule functions in cellular processes.

### Technical Limitations and Future Directions

Our Bio-ID system enables us to identify proteins that are specifically located within the microtubule lumen, overcoming the limitation of previous studies that could not distinguish between intraluminal and extraluminal proteins. The specificity and efficiency of our system in identifying MIPs (e.g., JPT2, CSPP1, etc.) represents a major advance in the field of microtubule research. However, while our approach successfully captured key MIPs in HeLa cells, it may have limitations in detecting dynamic protein interactions or in different cellular contexts. In particular, the space within the microtubule lumen is small, and the steric hindrance of BirA itself may affect the microtubule lumen proteins. Future studies should aim to improve this system to capture a wider range of MIPs in various cell types and physiological conditions.

### Role of MIPs in Paclitaxel Sensitivity and Therapeutic Potential

MIPs play a critical role in maintaining the structural and functional integrity of the microtubule lumen, influencing various cellular processes such as microtubule dynamics, stability, and drug interactions (68, 69). Our study identified 29 Paclitaxel-sensitive MIPs, including JPT2, CSPP1and JPT1, which localize within the microtubule lumen and modulate the response of cells to chemotherapy drugs like Paclitaxel. This discovery reveals that the microtubule lumen is not a passive compartment but a dynamic environment where MIPs can regulate microtubule function and influence the binding and efficacy of chemotherapeutic agents.

Paclitaxel stabilizes microtubules, arresting cell cycle and inducing apoptosis in cancer cells (67). However, resistance to Paclitaxel is a major challenge (70, 71). Our research suggests that MIPs may impact the sensitivity of cancer cells to Paclitaxel by competing for binding sites or altering microtubule structure. This modulation could affect the effectiveness of Paclitaxel’s therapeutic effects.

Moreover, different MIPs appear to have varying impacts on Paclitaxel sensitivity. Proteins such as CSPP1 is known to be involved in microtubule stabilization and organization, and their presence within the microtubule lumen may influence how Paclitaxel interacts with microtubules. The identification of these Paclitaxel-sensitive MIPs suggests that they may serve as potential targets for overcoming drug resistance. By targeting MIPs that modulate microtubule stability, it may be possible to enhance Paclitaxel’s efficacy in cancer cells that have developed resistance to treatment.

Our study suggests that MIPs are crucial in regulating microtubule dynamics and drug interactions, such as with Paclitaxel, making them promising therapeutic targets in cancer treatment. Targeting MIPs could enhance cancer cell sensitivity to drugs like Paclitaxel, potentially reducing side effects and enabling new combination therapies. A deeper understanding of MIPs may lead to the development of novel therapeutics that target the microtubule lumen, offering more effective treatments for drug-resistant tumors. Further research, including structural studies using cryo-electron microscopy as well as advanced technologies such as artificial intelligence, is needed to uncover the precise interactions between MIPs, microtubules, and chemotherapeutic drugs, paving the way for improved drug efficacy and the overcoming of drug resistance.

## Materials and Methods

### Cell culture

HeLa cells and AGS cells were maintained in DMEM/F12 (Wisent, CA) supplemented with 10% FBS (VivaCell, CN), 1% streptomycin and penicillin (Wisent, CA) at 37 °C. HEK293T cells were maintained in DMEM (Wisent, CA) supplemented with 10% FBS (VivaCell, CN), 1% streptomycin and penicillin (Wisent, CA) at 37 °C. MKN7 cells and MKN74 were maintained in RPMI-1640 (Gibco, USA) supplemented with 10% FBS (VivaCell, CN), 1% streptomycin and penicillin (Wisent, CA) at 37 °C. HeLa cells and HEK293T cells were obtained from Riken (JPN), AGS cells were obtained from BOSTER (CN), MKN74 cells and MKN7 cells were obtained from PROCELL (CN).

### Plasmids

All the sgRNAs oligonucleotides were annealed and ligated in the PX459 vector (26, 27). For constructed knock-in donor plasmid, TUBB4B homologous arm sequences were amplified from human genome DNA, and insert amplified from BirA* plasmid, which was a gift from Kyle Roux (Addgene plasmid # 36047; http://n2t.net/addgene:36047), three donor DNA fragments cloned into pUC19 vector. CSPP1 (XM_005251305.5), MAP6 (NM_033063.2), MATCAP1 (NM_001369686.1), and MEC17/ATAT1 (NM_001413067.1) were amplified from human cDNA and cloned into the pCDNA3.1-Kozak-Flag-mEGFP-N (homemade), pCDNA3.1-Kozak-HA-N (homemade) and pCDNA3.1-Flag-C vectors. MATCAP1 next subclone to pCDNA3.1-Kozak-Flag-N (homemade) vector for immunoprecipitation experiment. MEC17 subcloned to pCDNA3.1-mEGFP-C (homemade) vector for immunofluorescence staining. JPT2 (NM_144570.3) was amplified from human cDNA and cloned into the pCDNA3.1-Kozak-Flag-copGFP-N (homemade) vector, next for purified JPT2 or mEGFP-JPT2 proteins, JPT2 was subcloned into pHUE-10His-Ub-N (donated by Dr. Liu) or pCDNA3.1-Kozak-Strep tag 2-mEGFP-N (homemade) vectors. To generate JPT2 stable overexpression cells, subcloned copGFP-JPT2 into pLVX-IRES-Puro (donated by Dr. Guo) vector. To construct JPT2 knock-in donor plasmid, homologous arm sequences were amplified from human genome DNA, and copGFP fragment was amplified from pCDNA3.1-Kozak-Flag-copGFP-N, three donor DNA fragments cloned into pUC19 vector. For confirmed JPT2 microtubule binding motif, multiple JPT2 truncate and deletion mutants were subcloned by pCDNA3.1-Kozak-Flag-copGFP-JPT2, and all the mutants were cloned into pCDNA3.1-Kozak-Flag-copGFP-N vector. For JPT2 SMLM imaging, JPT2 was amplified from pCDNA3.1-Kozak-Flag-copGFP-JPT2 and cloned into the pCDNA3.1-Kozak-mEGFP-N vector. SIRT2 (NM_012237.4) and HDAC6 (NM_006044.4) were amplified from human cDNA and cloned into the pCDNA3.1-Kozak-mEGFP-N vector. To detect JPT2 relation with MEC17, MEC17 was amplified from pCDNA3.1-MEC17-Flag and cloned into the pCDNA3.1-TDtomato-Flag-C-IRES-Flag-copGFP (homemade) vector; and JPT2 fragment was amplified from pLVX-CopGFP-JPT2-IRES-Puro vector, then cloned into the pCDNA3.1-TDtomato-Flag-MEC17-IRES-Flag-copGFP-N vector. To generate JPT2 and JPT2 same dose stable overexpression knock-in cells, GAPDH homologous arm sequences were amplified from human genome DNA, and HA-BirA* fragment was amplified from pUC-19-TUBB4B-BirA*-donor plasmid, then JPT2 or JPT2-mutant cloned from pCDNA3.1-Kozak-Flag-CopGFP-JPT2 or pCDNA3.1-Kozak-Flag-CopGFP-JPT2-mutant vectors, three or four donor DNA fragments cloned into pUC19 vector. JPT1 (NM_016185.4) was amplified from human cDNA and cloned into the pCDNA3.1-Kozak-Flag-mEGFP-N and pCDNA3.1-Kozak-HA vector. Next JPT1 subcloned to pCDNA3.1-Kozak-Flag-copGFP-N vector for immunofluorescence staining. pCDNA3.1-Kozak-copGFP-IRES-EB3-mScarlet and pCDNA3.1-Kozak-copGFP-JPT2-IRES-EB3-mScarlet cloned form copGFP-JPT2 and EB3 (purchased form Evrogen company) vectors, for microtubule dynamics experiment. Jupiter C terminal 17 a.a. was amplified from fruitfly cDNA and cloned to the pCDNA3.1-Kozak-Flag-copGFP-N vector.

### Generating knock-in HeLa cells

To generate an endogenous HA-BirA*-TUBB4B cell line, we used homology-directed repair and CRISPR-Cas9. TUBB4B sgRNA (5′-GTGCACGATTTCCCTCATGA-3′), to tag TUBB4B locus with HA-BirA* at the N terminus, homologous arm length 800bp.1.5μg donor plasmid was co-transfected with 1 μg PX459 (Cas9-guide RNA) into HeLa cells grown on six-well plates. 24 h post-transfection, cells were first selected with 2 μg/mL puromycin for 2 days, followed by serial dilution in 96-well plates. Positive clone cells were screened by immunofluorescence.

To generate an endogenous TUBB4B-BirA*-HA cell line, we used homology-directed repair and CRISPR-Cas9. TUBB4B sgRNA (5′-GCCTAGAGCCTTCAGTCACT-3′), to tag TUBB4B locus with BirA*-HA at the C terminus, homologous arm length 800 bp.1.5 μg donor plasmid was co-transfected with 1μg PX459 (Cas9-guide RNA) into HeLa cells grown on six-well plates. 24 h post-transfection, cells were first selected with 2 μg/mL puromycin for 2 days, followed by serial dilution in 96-well plates. Positive clone cells were screened by immunofluorescence.

To generate an endogenous CopGFP-JPT2 cell line, we used homology-directed repair and CRISPR-Cas9. JPT2 sgRNA (5′-TTCCAGGTCCCGGATAGCGA-3′), to tag JPT2 locus with CopGFP at the N terminus, homologous arm length 800 bp.1.5 μg donor plasmid was co-transfected with 1 μg PX459 (Cas9-guide RNA) into HeLa cells grown on six-well plates. 24 h post-transfection, cells were first selected with 2 μg/mL puromycin for 2 days, followed by serial dilution in 96-well plates. Positive clone cells were screened by immunofluorescence.

To generate an endogenous GAPDH-P2A-BirA*-HA, GAPDH-P2A-BirA*-HA-JPT2-mutant and GAPDH-P2A-BirA*-HA-JPT2 cells, we used homology-directed repair and CRISPR-Cas9. GAPDH sgRNA (5′-AGCCCCAGCAAGAGCACAAG-3′), to tag GAPDH locus with P2A-BirA*-HA, P2A-BirA*-HA-JPT2-mutant and P2A-BirA*-HA-JPT2 at the C terminus, homologous arm length 800 bp.1.5 μg donor plasmid was co-transfected with 1 μg PX459 (Cas9-guide RNA) into HeLa cells grown on six-well plates. 24 h post-transfection, cells were first selected with 2 μg/mL puromycin for 2 days, followed by serial dilution in 96-well plates. Positive clone cells were screened by immunofluorescence (61).

### Generating knock-out HeLa cells

To generate an JPT2 knock-out cell line, we used CRISPR-Cas9, JPT2 sgRNA (5′-ATCCTATTAGGCCTGCTGGA -3′). 2.5 μg PX459 (Cas9-guide RNA) into HeLa cells grown on six-well plates. 24 h post-transfection, cells were first selected with 2 μg/mL puromycin for 2 days, followed by serial dilution in 96-well plates. Positive clone cells were screened by western blotting.

To generate a MEC17-JPT2 double knock out cell line, we used CRISPR-Cas9, MEC17 sgRNA (5′-ATGACCATATATATGAACT -3′). 2.5 μg PX459 (Cas9-guide RNA) into JPT2-KO cells grown on six-well plates. 24 h post-transfection, cells were first selected with 2 μg/mL puromycin for 2 days, followed by serial dilution in 96-well plates. Positive clone cells were screened by western blotting and immunofluorescence staining.

### Generating JPT2 stable overexpression HeLa cells

JPT2 overexpression cell line was generated using lentiviral particles produced by transfecting pLVX-CopGFP-JPT2-IRES-Puro, psPAX2, and pMD2.G into HEK293T cells with Lipofectamine 2000. Cells were cultured for 3 days and collected medium then filtered through a 0.45-μm filter. Viruses were spined at 20,000 rpm for 2 h at 4 °C, removed supernatant then resuspended in 50μL PBS. HeLa cells were seeded in the 6-well plate, then added 2uL JPT2 viruses, and selected with puromycin (1 μg/mL), followed by serial dilution in 96-well plates, positive clone cells were screened by immunofluorescence.

### BioID and LC-MS/MS

BioID pull-down experiments for purifying biotin-labeled candidate singlet MIPs. HA-BirA*-TUBB4B knock-in cells, TUBB4B-BirA*-HA knock-in cells, and wild-type HeLa cells were seeded in two 15-cm dishes. Then, they were incubated for 12 hours in the medium supplemented with 50μM biotin, after washed PBS three times, harvested cells by lysis buffer (50 mM Tris-HCl, pH 7.4; 500 mM NaCl, 0.2% SDS, 1× protease inhibitor and 1 mM DTT), lysate on ice keep 30 min, then lysate was centrifuged at 50000 rpm for 45 min at 4 °C to remove the cellular debris. The supernatants were incubated with Dynabeads, overnight at 4 °C. Subsequently, the beads were washed three times at 25 °C in 1 M KCl and then washed thrice at 8 M uera with 50 mM NH₄HCO₃. After removing the supernatant completely, the beads with bound proteins were rinsed in 50 mM NH4HCO3. The proteins were reduced with 10 mM DTT at 37 °C for 1 hour and alkylated with 55 mM iodoacetamide at room temperature for 1 hour in the dark. On-bead trypsin digestion was performed at 37 °C overnight. The resultant tryptic peptides were desalted by StageTips and dried with a SpeedVac concentrator. MS analyses were performed as described in the “proteomic profiling” part. The database search was performed for all raw MS files using the software MaxQuant (version 1.6) (28).

### Bio-IP and western blotting

Overexpression of Flag-mEGFP-MAP6, Flag-MATCAP1, Flag-mEGFP-CSPP1, MEC17-Flag, Flag-CopGFP-JPT2 and Flag-CopGFP-JPT1 in HA-BirA*-TUBB4B knock-in cells with Lipofectamine 2000. Cells were incubated for 12 hours in the medium supplemented with 50 μM biotin (Sigma-Aldrich) when post-transfection 24 h, after washing PBS three times, harvested cells by lysis buffer (50 mM Tris-HCl, pH 7.4; 200 mM NaCl, 1 mM EDTA, 1% NP-40 and 1× protease inhibitor), the lysate was centrifuged at 50000 rpm for 45 min at 4 °C to remove the cellular debris. The supernatants were incubated with Flag antibody-conjugated beads or GFP antibody-conjugated beads, for 3 hours at 4 °C. Subsequently, the beads were washed three times at 25 °C in lysis buffer, removing the supernatant completely, the beads with bound proteins were rinsed in SDS-sample buffer (50 mM Tris-HCl, pH 7.4; 10% sodium dodecyl sulfate, 1% 2-Hydroxy-1-ethanethiol and 0.1% bromophenol blue) 10 min at 100 ℃. Western blotting detected the biotin modification level, then striped the PVDF membrane and detected the Flag or GFP signal in the same PVDF membrane.

### Immunofluorescence Staining

To immunostaining singlet MIPs, cells were permeabilized before the fixation. Cells were washed with an microtubule-stabilizing buffer (60 mM Pipes, 25 mM Hepes, 10 mM EGTA, 2 mM MgCl_2_, pH 6.9), and then permeabilized cells in microtubule-stabilizing buffer with 0.05 % Triton X-100 (added freshly) for 30 s. Removed buffer immediately and fixed with 3% PFA with 0.01% glutaraldehyde at 37 ℃ for 10 min, the background fluorescence of glutaraldehyde was quenched by incubating cells with 0.1% NaBH4 in PBS for 7 min at room temperature. Then washed three times for 5 min with PBST (0.1% Triton X-100 in PBS). After blocking with 3% BSA in PBST for 30 minutes at room temperature, the cells were incubated with primary antibodies (GFP fusion proteins relied only on their own fluorescence, no antibody) at room temperature for 1 hour and then washed with PBST three times for 5 min. Subsequently incubated with secondary antibody for 1 h, and then washed with PBST three times for 5 min. Finally, the cells were mounted on slides with FluorSave reagent (Millipore).

### Protein purification

For the purification of mEGFP-JPT2, HEK293T cells were transiently transfected with polyethyleneimine (Linear, MW40000; YEASEN) with StrepII-mEGFP-JPT2 or StrepII-JPT2 plasmids. The cells were harvested 36 hours after transfection. Cells from two 15-cm dishes were lysed in 3 mL lysis buffer (50 mM HEPES, 300mM NaCl,1mM MgCl_2_, 1 mM DTT, 0.5% Triton X-100, pH 7.4) supplemented with protease inhibitors (MEI5BIO) on ice for 30 min. The lysate was centrifuged at 50000 rpm for 45 min at 4 °C to remove the cellular debris. The supernatant was incubated with 100 µL StrepTactin beads (GE Healthcare) for 2 h. Wash the beads three times with 1 mL wash buffer 1 (50 mM HEPES, 300 mM NaCl,1 mM MgCl_2_, 1 mM DTT, 0.5% Triton X-100, pH 7.4), Wash the beads three times with 1 mL wash buffer 2 (50 mM HEPES, 1 M NaCl, 0.5% Triton X-100, pH 7.4.), Wash the beads twice with 1 mL wash buffer 3 (50 mM HEPES, 150 mM NaCl, pH 7.4). The protein was eluted in 200 μL elution buffer (wash buffer 3 supplemented with 2.5 mM d-Desthiobiotin [Sigma-Aldrich]).

For the purification of the JPT2 protein, an Escherichia coli vector (pHU-His-UB) was constructed for the expression of JPT2 proteins as histidine-tagged ubiquitin fusions, and a histidine-tagged deubiquitylating enzyme was used to cleave these fusions, as reported previously (72). Briefly, the plasmid was incubated in a 37 ℃ shaker until OD600 reached 0.6. Recombinant proteins were expressed in Escherichia coli BL21 (DE3) host cells at 18℃ for 16 h in the presence of 0.4 mM IPTG. The cells were then collected by centrifugation. Cells pellets were resuspended in lysis buffer (20 mM Tris, pH 8.0, 150 mM NaCl), supplemented with protease inhibitor cocktail (mei5bio), sonicated to homogenize samples, then incubated 30 min after the addition of 1% Triton X-100. The mixture was pelleted by centrifugation at 260,000 g for 30 min at 4 ℃. Proteins were purified by Ni^2+^-Sepharose 6 Fast Flow (GE Healthcare) according to the manufacturer’s instructions, followed by size-exclusion chromatography (Superdex-75 10/300, GE Healthcare).

All purified proteins were snap-frozen in liquid nitrogen and stored at -80 ℃.

### Tubulin purification and in vitro microtubule dynamic assay

Crude tubulin was obtained from the pig brain by double cycles of polymerization and depolymerization, as previously described (73). Tubulin was further purified using a TOG-based affinity column (74). Tubulin was labeled with biotin (Thermo Fisher Scientific, 20217), TAMRA (Thermo Fisher Scientific, C1171), and Alexa Fluor 647 (Thermo Fisher Scientific, A20106) using NHS esters, following standard protocols (75).

The microtubule dynamic assay was performed as described previously (76). Short GMPCPP-stabilized microtubules (5% Alexa Fluor 647 labeled and 20% biotin-labeled) were immobilized on the surface of a cover glass coated with a biotin-binding protein (Thermo Fisher Scientific, 31000). Free tubulin dimer (13% TAMRA labeled) was then added into the flow cell in imaging buffer (BRB80 supplemented with 2 mM GTP, 50 mM KCl, 0.15% sodium carboxymethylcellulose, 80 mM D-glucose, 0.4 mg/mL glucose oxidase, 0.2 mg/mL catalase, 0.8 mg/mL casein, 1% β-mercaptoethanol, 0.001% Tween 20). The growth of microtubules was recorded using a total internal reflection (TIRF) microscope (Olympus) equipped with an Andor 897 Ultra EMCCD camera (Andor, Belfast, UK) using a 100 × TIRF objective (NA 1.49; Olympus). To record microtubule dynamics, images were taken every second with a 0.1 s exposure.

### Microtubule co-sedimentation assay

To detect JPT2 interaction with microtubules, GMPCPP-stabilized microtubules were incubated with JPT2 or BSA in the BRB80 buffer. Tubulin was assembled into GMPCPP microtubules in a reaction buffer (1 mM GMPCPP, 4 mM MgCl_2_, 8 μM tubulin in BRB80) at 37 ℃ overnight, Subsequently, 300 μL of BRB80 buffer was added, followed by centrifugation at 18,000 g for 20 min at 37 ℃. After removing the supernatant, the pellet was resuspended in the BRB80 buffer.

Proteins at a concentration of 1 μM were incubated with microtubules at various concentrations for 30 minutes at room temperature, followed by centrifugation at 18,000 g for 20 minutes at 37 ℃. After collecting the supernatant, the pellet was resuspended in BRB80 buffer equal to the volume of the supernatant. Subsequently, the samples were boiled with the SDS-PAGE sample buffer. All the samples were analyzed using SDS–PAGE. For analysis, the intensities of the tubulin and JPT2 or BSA protein bands were quantified using ImageJ. Y represents the fraction of JPT2 or BSA proteins found in the pellet.

### Rose-Z Imaging

Cells were permeabilized before the fixation for singlet MIPs 3D SMLM imaging. Cells were washed with microtubule-stabilizing buffer (60 mM Pipes, 25 mM Hepes, 10 mM EGTA, 2 mM MgCl_2_, pH 6.9), and then permeabilized cells in microtubule-stabilizing buffer with 0.05% Triton X-100 (added freshly) for 30 s. Removed buffer immediately and fixed with 3% PFA with 0.01% glutaraldehyde at 37 ℃ for 10 min. The background fluorescence of glutaraldehyde was quenched by incubating cells with 0.1% NaBH4 in PBS for 7 min at room temperature. Then, cells were washed with PBS three times and permeabilized with 0.2% Triton X-100 in PBS for 15 min.

Subsequently, cells were incubated for 90 min blocking with PBS containing 10% normal goat serum (NGS) and 0.05% Triton X-100 at room temperature, immunostained with primary antibodies (diluted with PBS containing 5% NGS and 0.05% Triton X-100) for 1 h, washed with washing buffer (1% NGS and 0.05% Triton X-100 in PBS) five times (15 min per wash), and incubated with secondary antibodies for 60 min. Cells were washed with PBS for 5 min after being washed with washing buffer. PBS containing 2% paraformaldehyde and 0.1% glutaraldehyde was used to post-fix cells for 10 min. Cells were then washed with PBS three times (5 min per wash). Finally, cells were washed with ddH_2_O twice (3 min per wash) and saved with ddH_2_O at 4 °C.

SMLM imaging and data analysis: The interferometric SMLM imaging was performed with a custom-built super-resolution microscope on an Olympus IX71 inverted microscope and a 100 × 1.5 NA oil-immersion objective. The microscope utilized interferometric fringes to achieve an ultra-high z-axis resolution, which is similar to that reported for lateral resolution improvement (50). A 639-nm laser was used for the excitation of Alexa Fluor 647 and CF 660C, and a 405-nm laser was used for activation and control of the molecule intensity during image acquisition. Images were acquired on an Andor iXon DU-897 EMCCD camera, and the pixel size was 150 nm with a field of view of ∼ 30 μm × 30 μm. An active drift correction method was used to correct mechanical drift during image acquisition. For SMLM data acquisition, an exposure time of 50 ms, an EM gain of 20, and an illumination power intensity of ∼4 kW/cm^2^ were used. A total of 400000 frames were acquired for the super-resolution image reconstruction. During imaging, the 405-nm laser was used to control the molecule density. The sample was sealed between two coverslips to create a sandwich structure for imaging, with imaging buffer, i.e. 10% glucose, oxygen removed, GLOX (0.6 mg/mL glucose oxidase and 0.06 mg/mL catalase dissolved in Tris-HCl buffer), and 143 mM 2-mercaptoethanol in PBS, filled in the chamber to enable blinking of the dyes during imaging. Sample drift during acquisition was calculated by reconstructing STORM images from subsets of frames and correlating images to a reference frame.

### Immunoprecipitation (IP)

Overexpression EGFP-HDAC6 and EGFP-SIRT2 with HA-BirA*-JPT2 in HeLa cells by Lipofectamine 2000. The transfected cells were harvested by cold lysis buffer (20 mM Tris-HCl pH7.4, 150 mM NaCl, 1 mM EDTA and 1% NP-40) containing protease inhibitor cocktail on ice for 30 min, then centrifuging to remove insoluble fraction, GFP antibody-conjugated beads were added into supernatant for 2 h to capture GFP fusion proteins. The beads were subsequently washed three times in the wash buffer (20 mM Tris-HCl pH7.4, 150 mM NaCl, 1 mM EDTA and 0.1% NP-40). Removing the supernatant completely, the beads with bound proteins were rinsed in SDS-sample buffer for 10 min at 100 ℃. Finally, Western blotting detected candidate proteins.

### EB3 tracking

For EB3 time-lapse imaging, HeLa cells were seeded in a 35 mm glass bottom dish, then cells were transiently transfected with CopGFP-IRES-EB3-mScarlet or CopGFP-JPT2-IRES-EB3-mScarlet by Lipofectamine 2000. Post-transfection 18 h, the dishes were placed in a sealed chamber at 37 °C with 5% CO_2_, images of living cells were taken with a TRIF-SIM system, the system constructed by the Institute of Biophysics and applied as described (77), interval time 1 s, total 60 s. MATLAB software u-track was used to analyze EB3 features (78).

### siRNA Knock down

For siRNA knock down experiments, cells were seeded in 6-well plates with a complete growth medium. Next, the cells were transfected siRNAs for 6 h by Lipofectamine 2000. Post-transfection 72 h, collected cells for Western blotting or CCK-8 assay. 50 nM siRNA oligonucleotides transfected in each well, and standard negative siRNA as control siRNA.

siRNA oligonucleotides were used in this publication:

JPT2 5′-GCAUCUCCUUCUACUAAGATT-3′;

JPT1 5′-GAGACUUCUUAGAUCUGAATT-3′;

CSPP1 5′-GCUGAAAGAUAGAGAUUCA-3′

control siRNA 5′-UUCUCCGAACGUGUCACGUTT -3′.

### Cell viability assay

To detect cells in Paclitaxel cytotoxicity experiments, cells were treated with different concentrations of Paclitaxel ranging from 0 nM to 1000 nM for 72 h. 5000 cells were seeded in 96-well-plate each well, five parallel compound holes. Before detection, the medium was removed and cells were washed once with PBS. Fresh medium containing CCK8 reagent (MCE). After incubated at 37 °C for 2 h, the A450 values were read by the microplate reader (EnSpire 2300, PerkinElmer).

### Antibodies

α-tubulin antibody (Mouse, DM1A, IF 1:1000, Sigma; Rabbit, ER130905, WB 1:2000, HUABIO), Ace-tubulin antibody (Mouse, 6-11B-1, IF:1:1000, WB:1:20000, Sigma), EB1 antibody (Mouse, 610535, IF 1:200, BD Biosciences), Flag antibody (Mouse, M2, IF 1:8000, Sigma; Rabbit, F7425, WB 1:10000, Sigma), HA antibody (Rat, 3F10, IF 1:500, Roche; Mouse, WB 1:2000, Covance), GFP antibody (Rabbit, 598, IF 1:1000, MBL; Mouse, 66002-1, WB 1:3000, Proteintech), JPT2 antibody (Rabbit, HPA041908, WB 1:2000, ATLAS Antibodies), JPT1 antibody (Rabbit, FNab03940, WB 1:1000, Finetest), CSPP1 antibody (Rabbit, 11931-1, WB 1:1000, Proteintech).

### Patient Survival Analysis

We downloaded mRNA expression z-scores relative to normal samples (log RNA Seq V2 RSEM) data from the TCGA dataset. Disease-specific survival was estimated for each gene expression cohort (high versus low) using the Kaplan-Meier method. Cohort survival differences were assessed using the log-rank test. A low expresser was defined as JPT1 < 3.5, JPT2 < 2.0 and CSPP1 < 0 for the TCGA.

## Supplementary Figure Legends

**Supplementary Figure 1. Establishment of the TUBB4B-BirA* knock-in cell line.**

**(A)** Diagram of the TUBB4B-BirA*-HA-N knock-in cell line. **(B)** Immunostaining for TUBB4B (green) in TUBB4B-BirA*-N knock-in HeLa cells. **(C)** Western blotting detected the biotin modification level of MIPs and the expression level of TUBB4B in TUBB4B-BirA*-N knock-in cells, with GAPDH serving as a reference protein. Three signals were collected from the same PVDF membrane. **(D)** Diagram of the TUBB4B-BirA*-C knock-in cell line. **(E)** Immunofluorescence staining of TUBB4B (green) in TUBB4B-BirA*-C knock-in HeLa cells. **(F)** Western blotting detected the biotin modification level of outer MAPs and the expression level of TUBB4B in TUBB4B-BirA*-C knock-in cells, with GAPDH serving as a reference protein. Three signals were collected from the same PVDF membrane. **(G)** Immunostaining of MIPs (CSPP1, MAP6, MEC17) and the microtubule surface protein MATCAP1, with co-staining for α-tubulin in HeLa cells. **(H)** The histogram shows the molecular weight distribution of known MIPs.

Scale bars: 10 μm.

**Supplementary Figure 2. JPT2 binds to microtubules through its C-terminal domain and PPGGK*S motifs.**

**(A)** The Scheme of the domain organization of copGFP-JPT2 and the truncated mutants were used to determine the microtubule-binding domain of JPT2. **(B)** Immunostaining for α-tubulin (red) in copGFP-JPT2 and mutants overexpressing HeLa cells. Scale bars: 10 μm.

**Supplementary Figure 3. JPT2 is a microtubule-binding protein and regulates microtubule dynamics in cells.**

**(A)** Coomassie Brilliant Blue staining of a gel with JPT2 purified from *E.coli*. **(B)** Microtubule co-sedimentation assays were performed to detect the direct binding capacity of JPT2 protein to microtubules. BSA is used as a negative control. **(C)** The trend line of the co-sedimentation ratio of JPT2 protein with microtubules significantly increases compared to BSA as the microtubule concentration increases. **(D)** Trace line of EB3 in HeLa cells overexpressing copGFP-IRES-EB3-mScarlet and copGFP-JPT2-IRES-EB3-mScarlet respectively, scale bars: 10 μm. **(E)** Quantification of microtubule plus-ends growth rate and EBs lifetime in (D). For the control group, n = 30 cells; for the copGFP-JPT2 group, n = 38 cells. Values = means ± SEM. ****p < 0.0001, unpaired two-tailed Student’s t-test.

**Supplementary Figure 4. JPT2 inhibits the acetylation of microtubules within cells not by recruiting deacetylases.**

**(A)** Immunostaining for Ace-tubulin (Red) and copGFP (Green) in GFP-JPT2 stable expressing HeLa cells treated with DMSO or TSA. Enlarged microtubules in (A) used for line scan analyses reflect the co-localization of JPT2 and microtubules. JPT2 and acetylated microtubules are arranged in an interlaced pattern, but as the acetylation level of microtubules within the cell increases, the co-localization ratio of JPT2 with acetylated microtubules also increases, scale bars: 10 μm. **(B)** Schematic representation of the genomic DNA structure and editing region of JPT2. The sequencing result of the JPT2 KO cell line near the sgRNA sequence was shown aligned with the wild-type sequence. **(C)** Western blot analysis of microtubule acetylation levels in JPT2 KO and control HeLa cells, with α-tubulin serving as a reference protein. **(D)** Immunostaining for Ace-tubulin (Red) in JPT2 KO and WT HeLa cells, scale bars: 10 μm. **(E)** Quantification of Ace-tubulin fluorescence intensity per cell in control and JPT2 KO HeLa cells. For the WT group, n = 100 cells; For the JPT2 KO group, n = 118 cells. Values = means ± SEM. P = 0.1059, unpaired two-tailed Student’s t-test. **(F)** Co-immunoprecipitation assays were performed to detect the interaction between JPT2 and deacetylases HDAC6 and SIRT2. The experimental results showed that JPT2 did not interact with HADC6 or SIRT2. **(G)** Schematic representation of the genomic DNA structure and editing region of MEC17. The sequencing result of the JPT2-MEC17 dual KO cell line near the sgRNA sequence was shown aligned with the wild-type sequence. **(H)** Western blotting detected JPT2 and Ace-tubulin expressing levels in the JPT2-MEC17 dual KO cell line, α-tubulin as a reference protein.

**Supplementary Figure 5. The JPT2 family proteins and their homologs possess the same biological function.**

**(A)** Sequence alignment of JPT2, JPT1, and their homolog Jupiter. **(B)** Immunostaining for α-tubulin (Red) or Ace-tubulin (Red) in copGFP-Jupiter overexpressed HeLa cells, scale bars: 10 μm. **(C)** Immunostaining for Ace-tubulin (Red) in Flag-copGFP and copGFP-Jupiter-17 a.a. overexpressed HeLa cells, scale bars: 10 μm. **(D)** Quantification of Ace-tubulin fluorescence intensity per cell in copGFP and copGFP-Jupiter-17 a.a. overexpressed HeLa cells. For the copGFP group, n = 116 cells; for the copGFP-Jupiter-17 a.a. group, n = 151 cells. Values = means ± SEM. ****p < 0.0001, unpaired two-tailed Student’s t-test. **(E)** Western blot analysis of Ace-tubulin expressing levels in copGFP and copGFP-Jupiter-17 a.a. overexpressed HeLa cells, with α-tubulin serving as a reference protein.

**Supplementary Figure 6. Paclitaxel dissociates JPT2 but not MAP6 from microtubules.**

**(A)** Immunostaining for α-tubulin (red) in DMSO and Paclitaxel-treated copGFP-JPT2-178-190 a.a. expressing HeLa cells, scale bars: 10 μm. **(B)** Quantification of the copGFP-JPT2-178-190 a.a. intensity in DMSO and Paclitaxel-treated HeLa cells. For the DMSO group, n = 69 cells; For the taxol group, n = 113 cells. **(C)** Immunostaining for mEGFP-MAP6 (Green) and α-tubulin (red) in DMSO and Paclitaxel-treated HeLa cells, scale bars: 10 μm. **(D)** Quantification of the MAP6 intensity in DMSO and Paclitaxel-treated HeLa cells. The mean intensity of cells treated with Paclitaxel was normalized to the mean intensity in cells treated with DMSO. For the DMSO group, n = 67 cells; for the Taxol group, n = 91 cells.

Values = means ± SEM. ****p < 0.0001, p = 0.1253, unpaired two-tailed Student’s t-test.

**Supplementary Figure 7. Taxanes dissociate JPT2 from microtubules.**

**(A)** Immunostaining for EB1 (Gray) in DMSO and microtubule-stabilizing drugs-treated HeLa cells, Scale bars: 10 μm. **(B)** Quantification of the EB1 comets number per cell in DMSO and microtubule-stabilizing drugs-treated HeLa cells. For the DMSO group, n = 29 cells; Taxol group, n = 32 cells; Epothilone D group, n = 32 cells; TPI287 group, n = 40 cells; Noscapine group, n = 34 cells; TTI237 group, n = 34 cells. **(C)** Immunostaining for α-tubulin (red) in DMSO and Paclitaxel-treated copGFP-JPT2 stable expressing HeLa cells, scale bars: 10 μm. **(D)** Quantification of the JPT2 intensity in DMSO and microtubule-stabilizing drugs-treated HeLa cells. For the DMSO group, n = 51 cells; Taxol group, n = 53 cells; Epothilone D group, n = 53 cells; TPI287 group, n = 57 cells; Noscapine group, n = 49 cells; TTI237 group, n = 53 cells.

Values = means ± SEM. ****p < 0.0001, unpaired two-tailed Student’s t-test.

**Supplementary Figure 8. Paclitaxel dissociates CSPP1 and JPT1 from microtubules.**

**(A)** Time-lapse images of Paclitaxel-tracker (red) binding microtubule experiment in mEGFP-JPT1 and mEGFP-CSPP1 overexpressed HeLa cells. The right panel shows the quantification of the mean Docetaxel intensity in mEGFP-JPT1 and mEGFP-CSPP1 overexpressed Hela cells and control cells, scale bars: 10 μm. For the control group, n = 22 cells, for the JPT1-OE group, n = 17 cells. For the control group, n = 28 cells, for the CSPP1-OE group, n = 20 cells. The average mean intensity of overexpressing cells was normalized to the average mean intensity in control cells. **(B)** Survival curves of endometrial cancer with different expression levels of Paclitaxel-sensitive microtubule-inner protein mRNA following Paclitaxel treatment in TCGA database.

## Acknowledgements

We thank WX.Guo ( Institute of Genetics and Developmental Biology) for providing pLVX-IRES-Puro plasmid; CM.Liu( Institute of Genetics and Developmental Biology) for providing pHUE-10His-Ub-N plasmid; QF.Wu ( Institute of Genetics and Developmental Biology) for providing px459 plasmid. This work was funded by the National Natural Science Foundation of China(31930025,2021YFA0804802).

