## Supplemental Figures for "Systematic Identification of Microtubule Inner Proteins Reveals JPT2 as a Key Regulator of Lumen Microenvironment and Drug Sensitivity"

### Supplemental Figure 1

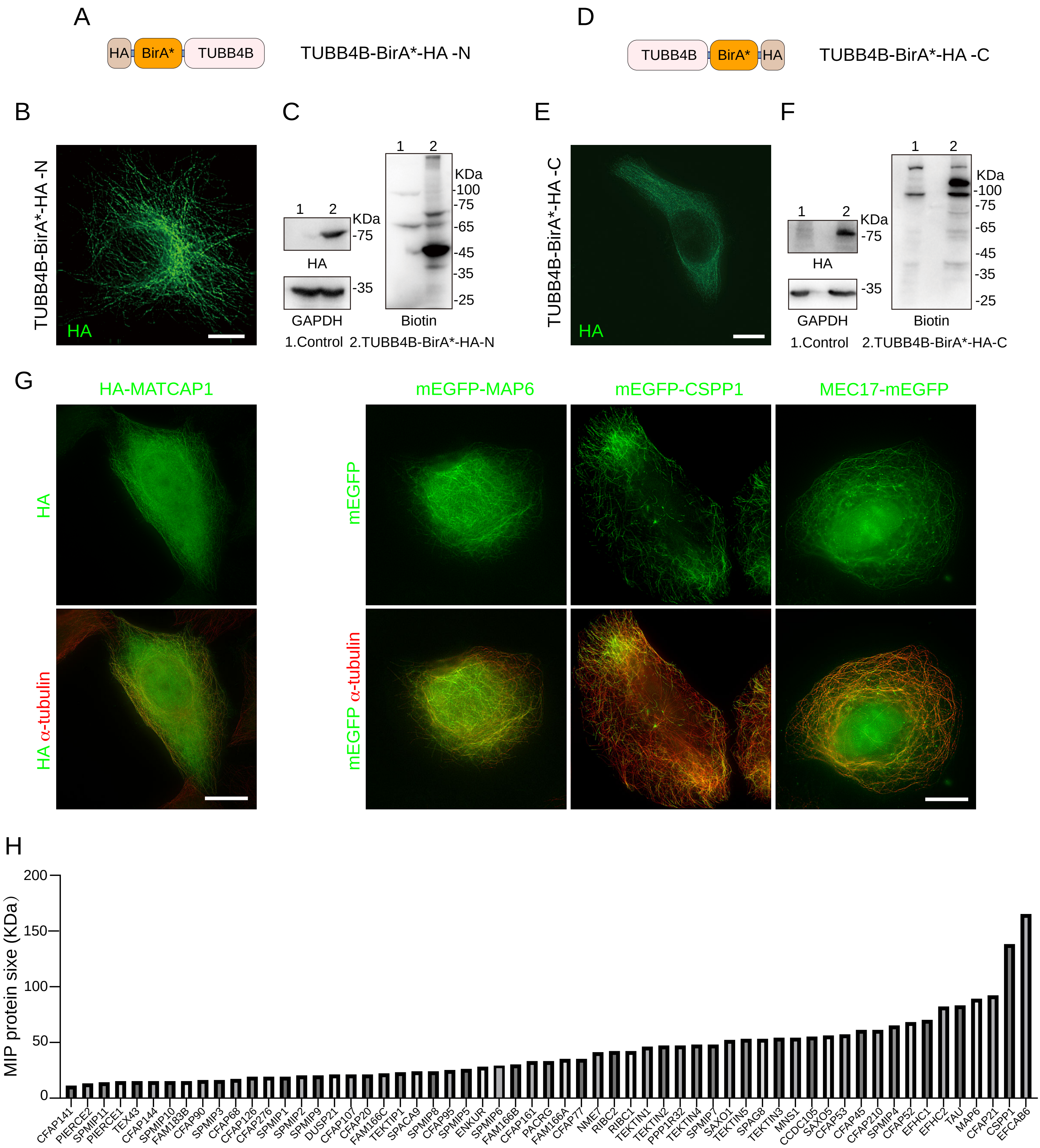

### Supplemental Figure 2

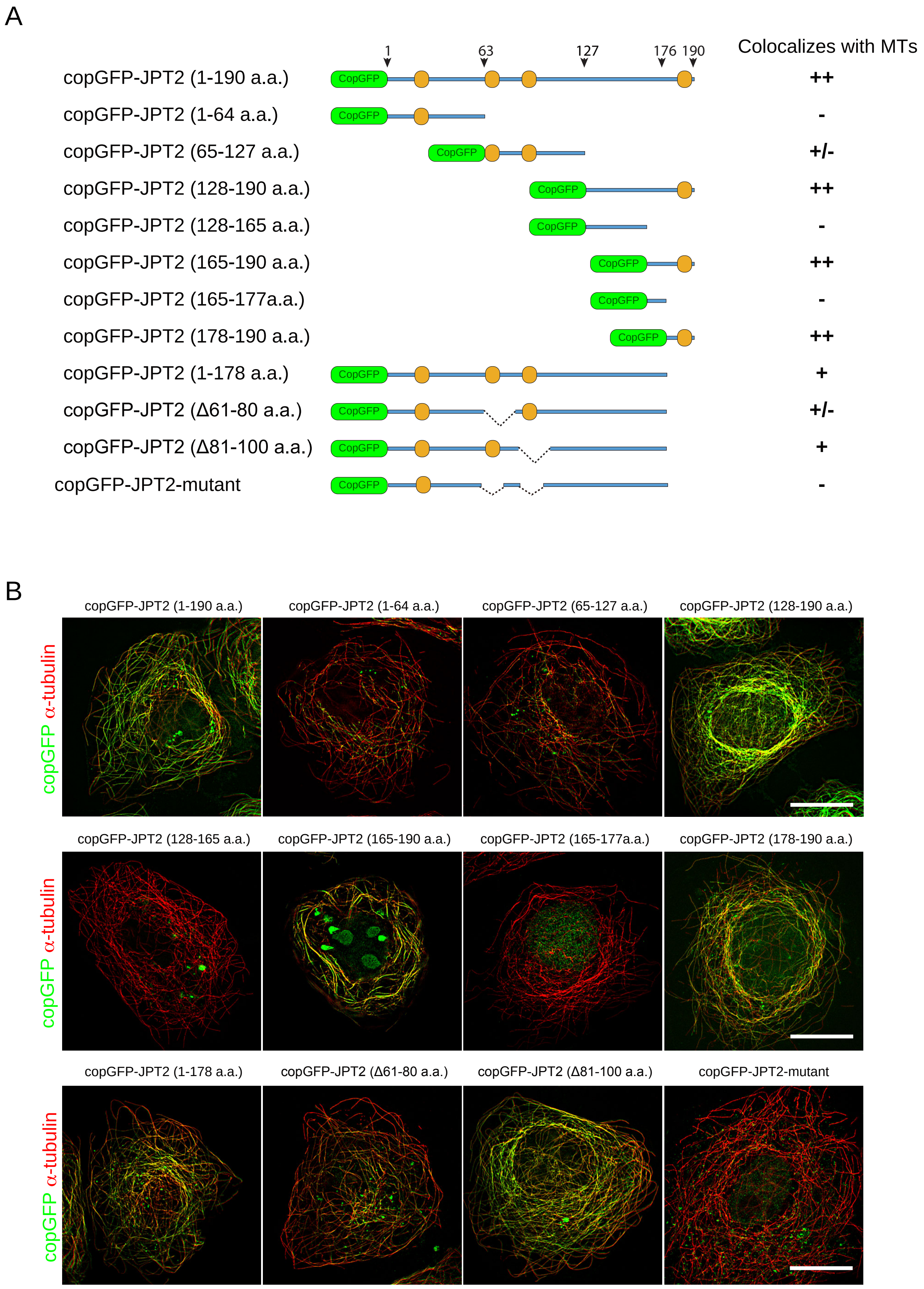

### Supplementary Figure 3

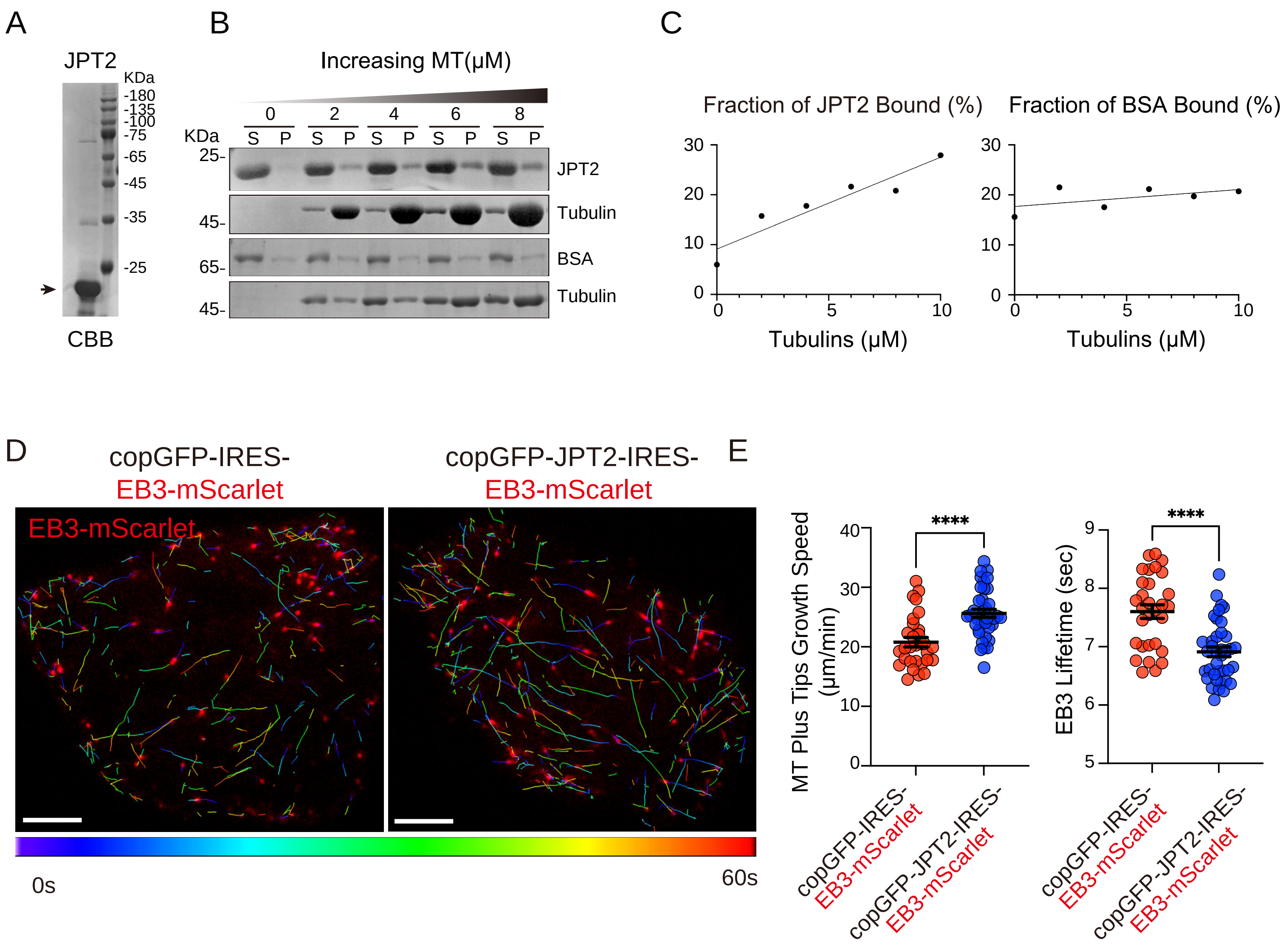

### Supplemental Figure 4

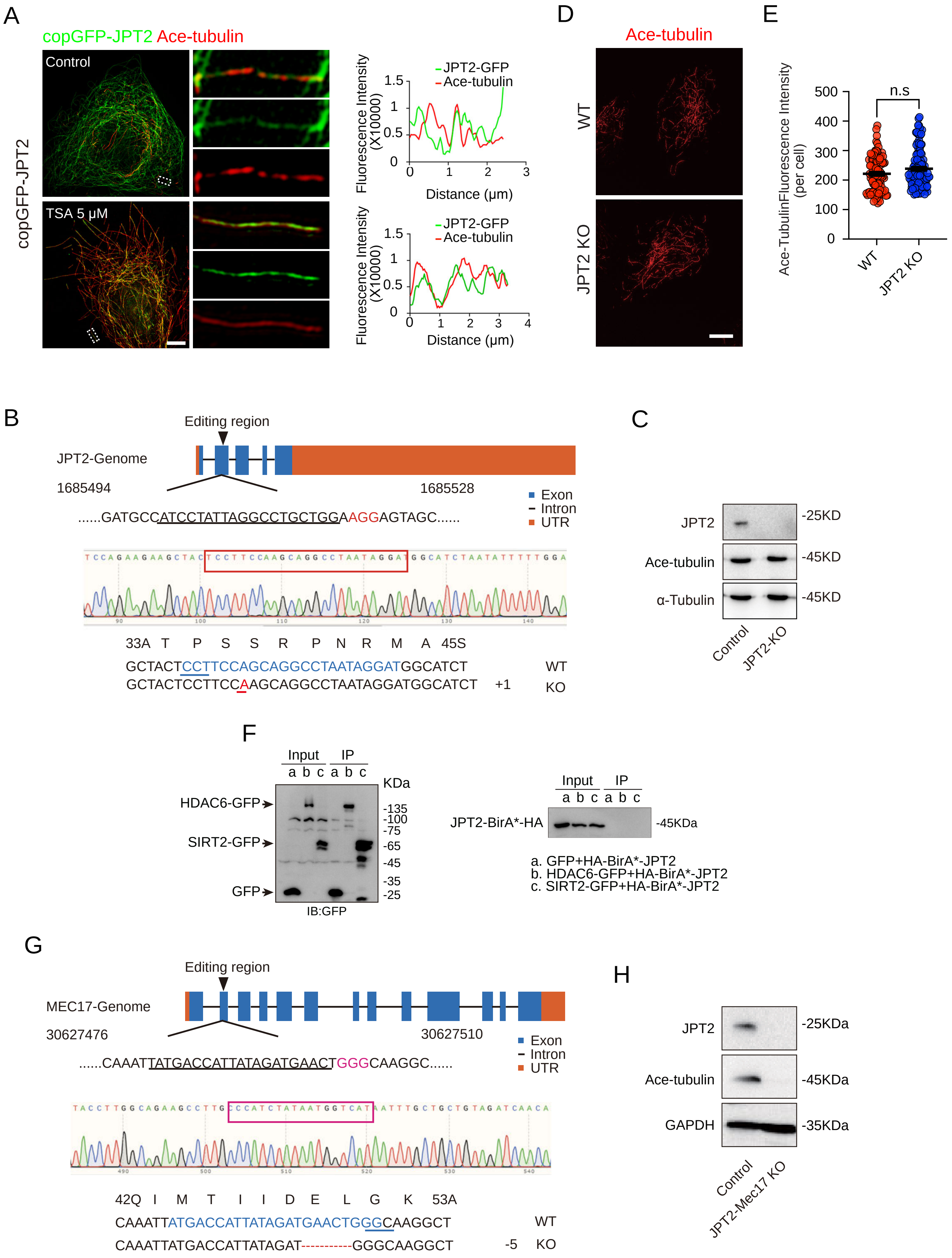

### Supplemental Figure 5

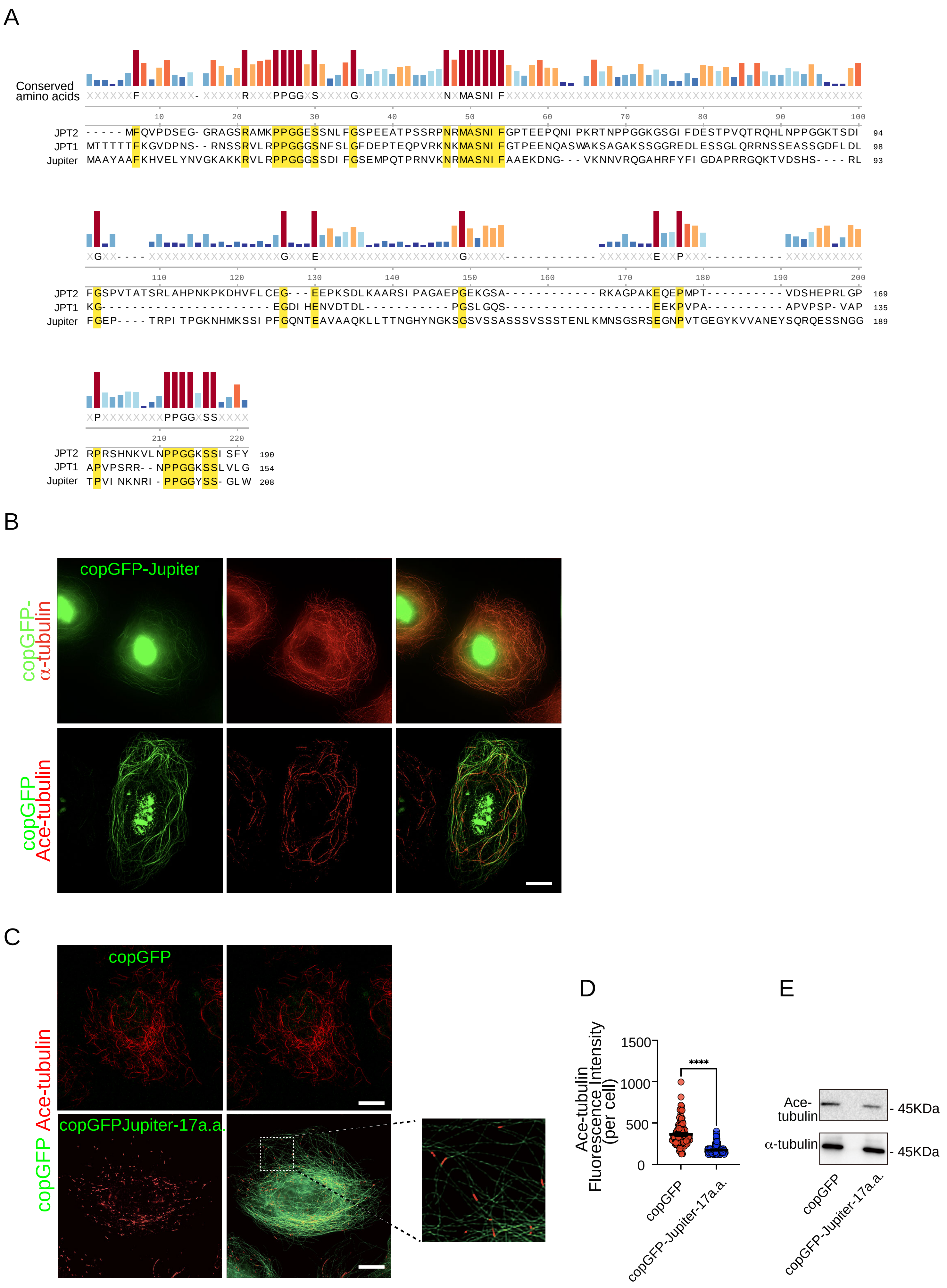

### Supplementary Figure 6

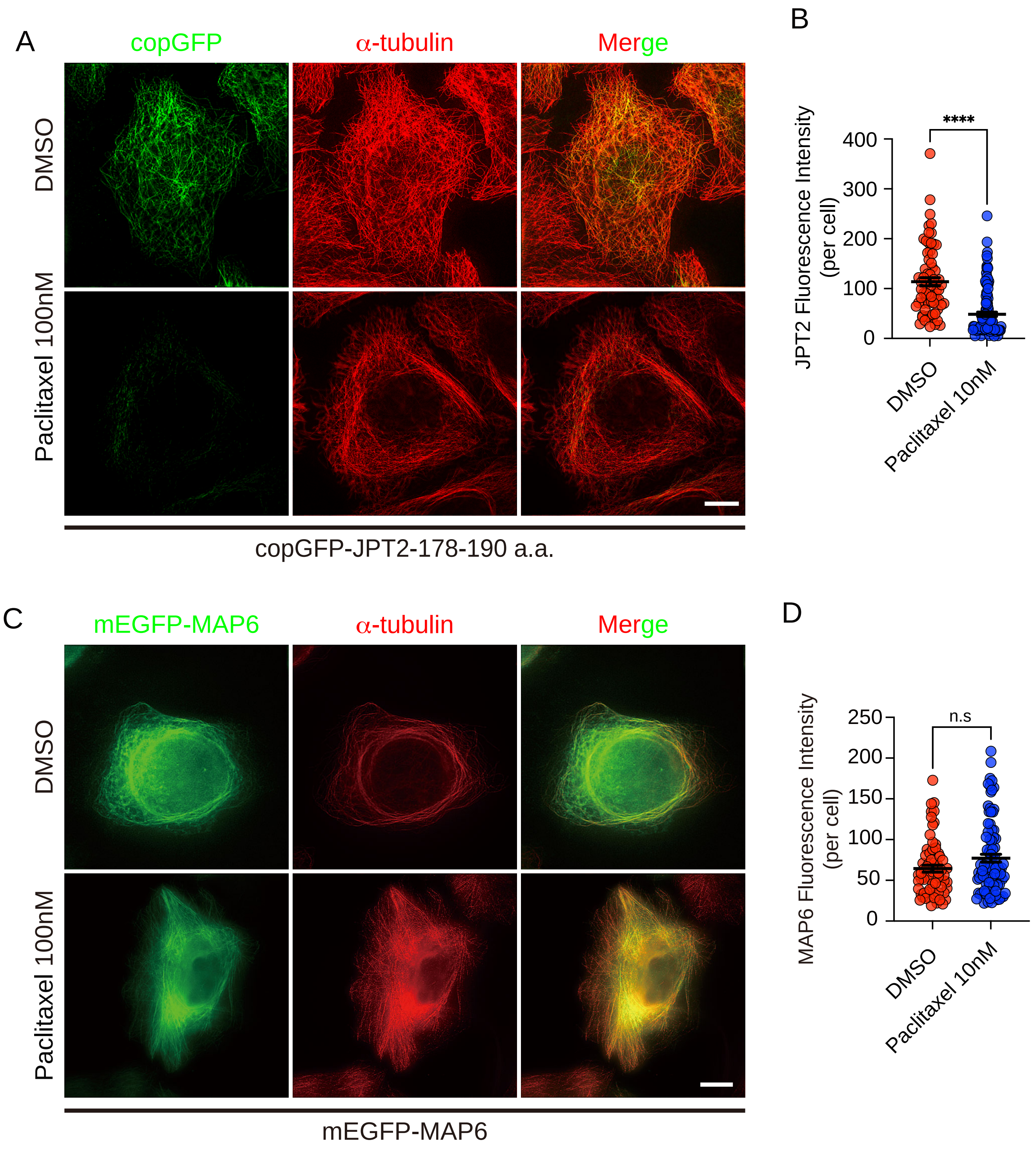

### Supplementary Figure 7

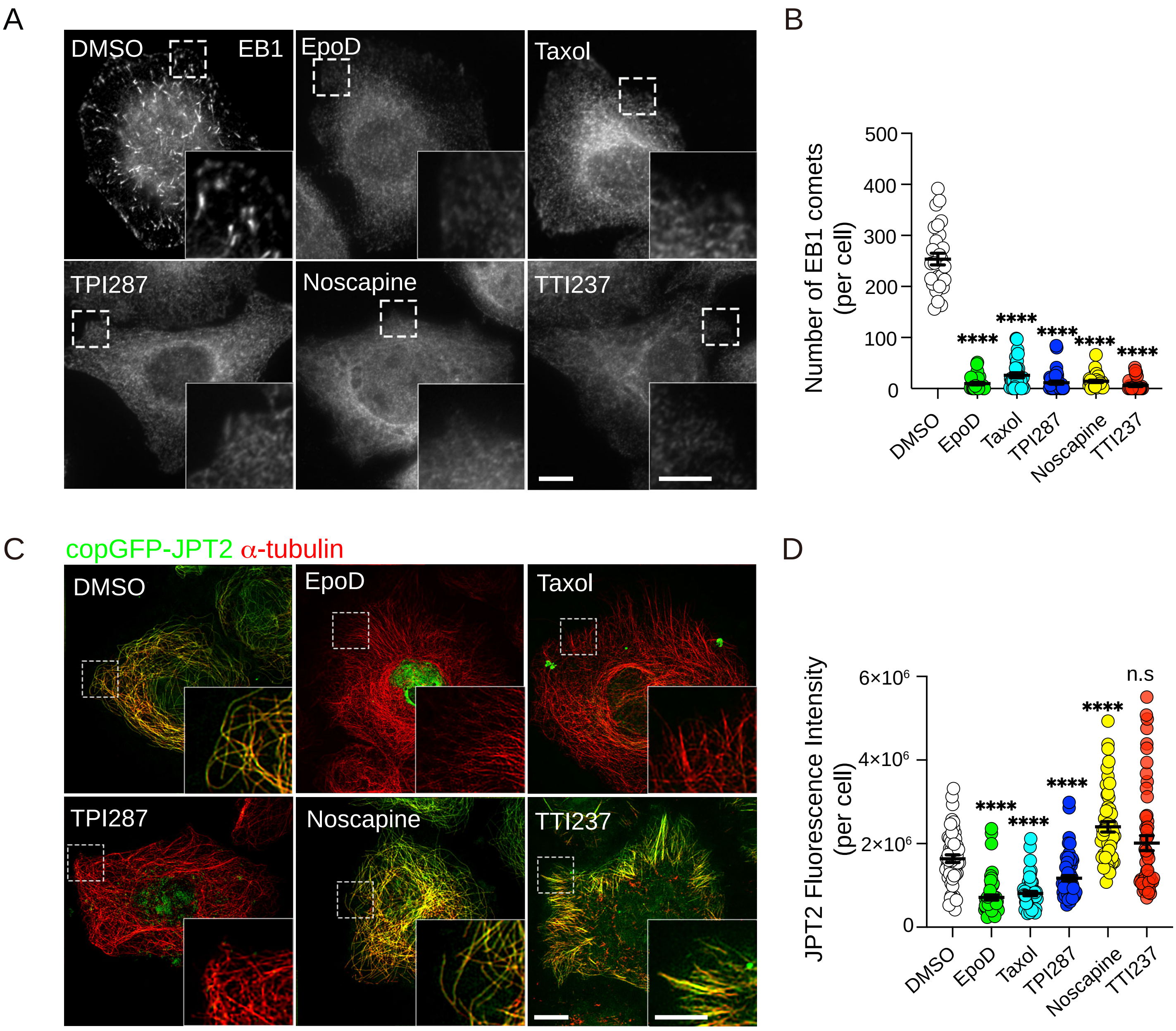


copGFP-JPT2 stably expressed HeLa cells

### Supplementary Figure 8

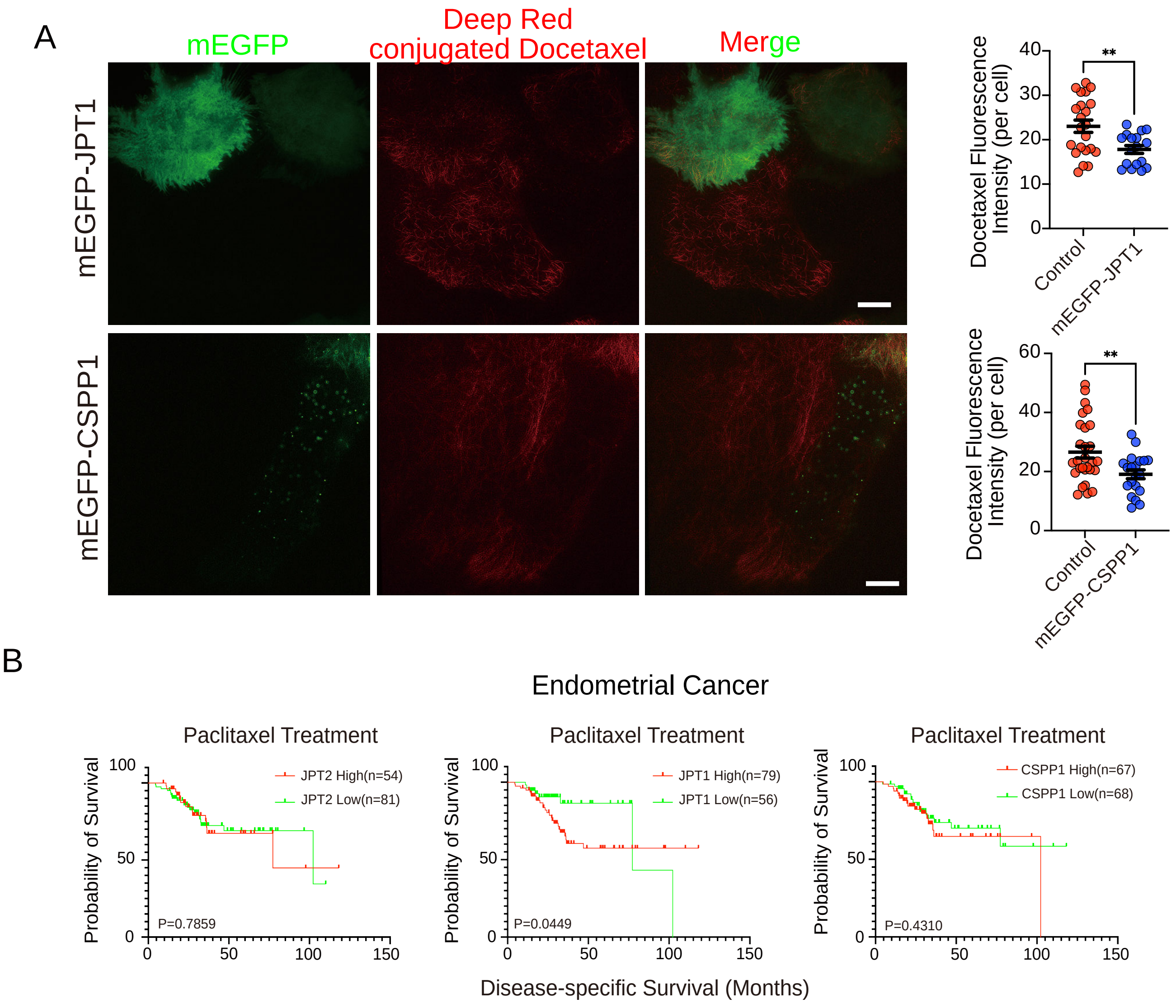
